# Recurrent mobilization of ancestral and novel variants of the chromosomal di-hydrofolate reductase gene drives the emergence of clinical resistance to trimethoprim

**DOI:** 10.1101/2020.07.31.230557

**Authors:** Miquel Sánchez-Osuna, Pilar Cortés, Montserrat Llagostera, Jordi Barbé, Ivan Erill

## Abstract

2.

Trimethoprim is a synthetic antibacterial agent that targets folate biosynthesis by competitively binding to the di-hydrofolate reductase enzyme (DHFR). Trimethoprim is often administered synergistically with sulfonamide, another chemotherapeutic agent targeting the di-hydro-pteroate synthase (DHPS) enzyme in the same pathway. Clinical resistance to both drugs is widespread and mediated by enzyme variants capable of performing their biological function without binding to these drugs. These mutant enzymes were assumed to have arisen after the discovery of these synthetic drugs, but recent work has shown that genes conferring resistance to sulfonamide were present in the bacterial pangenome millions of years ago. Here we apply phylogenetics and comparative genomics methods to study the largest family of mobile trimethoprim resistance genes (*dfrA*). We show that most of the *dfrA* genes identified to date map to two large clades that likely arose from independent mobilization events. In contrast to sulfonamide resistance (*sul*) genes, we find evidence of recurrent mobilization in *dfrA* genes. Phylogenetic evidence allows us to identify novel *dfrA* genes in the emerging pathogen *Acinetobacter baumannii*, and we confirm their resistance phenotype *in vitro*. We also identify a cluster of *dfrA* homologs in cryptic plasmid and phage genomes, but we show that these enzymes do not confer resistance to trimethoprim. Our methods also allow us to pinpoint the chromosomal origin of previously reported *dfrA* genes, and we show that many of these ancient chromosomal genes also confer resistance to trimethoprim. Our work reveals that trimethoprim resistance predated the clinical use of this chemotherapeutic agent, but that novel mutations have likely also arisen and become mobilized following its widespread use within and outside the clinic. This work hence confirms that resistance to novel drugs may already be present in the bacterial pangenome, and stresses the importance of rapid mobilization as a fundamental element in the emergence and global spread of resistance determinants.

**Impact statement:** Antibiotic resistance is a pressing and global phenomenon. It is well-established that resistance to conventional antibiotics emerged millions of years ago in either antibiotic-producing bacteria or their competitors. Resistance to synthetic chemotherapeutic agents cannot be explained by this paradigm, since these drugs are not naturally produced. Resistance is hence assumed to have evolved rapidly following the clinical introduction of these drugs. Recently we showed that resistance to one such drug, sulfonamide, evolved not recently, but millions of years ago, suggesting that the diversity of bacterial genomes may well contain genes conferring resistance to drugs yet to be developed. Here we analyze the origin of resistance to trimethoprim, another chemotherapeutic agent developed in the 1960’s. Using phylogenetic methods, we identify new variants of the trimethoprim resistance genes that had not previously been reported, and we trace the chromosomal origins for a number of already known resistance variants. Our results show that resistance to trimethoprim is very diverse and has originated both from recent mutations and from preexisting ancient variants. These results stress the importance of gene mobilization mechanisms as the main drivers of the current antibiotic resistance phenomenon.

**Data summary:**

- The scripts used for data collection and analysis can be obtained at the GitHub ErillLab repository (https://github.com/ErillLab/).
- The Bayesian phylogenetic tree can be visualitzed online on iTOL (https://itol.embl.de/tree/855674159451585133078) [1].
- The authors confirm all other supporting data has been provided within the article or through supplementary data files.

## 5. Introduction

Bacterial resistance to antibacterial agents remains an increasingly challenging and global problem in modern healthcare [2, 3]. Bacterial cells display a diverse array of mechanisms to cope with exposure to antibacterial compounds, including modification or overexpression of the antibacterial target, efflux or reduction of antibacterial uptake, and the use of alternate pathways [4]. Constant exposure to non-lethal concentrations of antibacterial agents may lead to the selection of partial resistance to antibiotics over relatively short timespans [5], and this evolution may be hastened by simultaneous exposure to multiple antibacterials [6]. However, the rapid proliferation of multidrug resistant nosocomial pathogens in the last fifty years has not been driven by the independent evolution of resistance traits, but through the extensive dissemination of mobile genetic elements carrying resistance genes [4, 7]. It is widely accepted that most genes conferring resistance to antibiotics present in pathogenic bacteria were obtained by successive lateral gene transfer of homologs that originally evolved in the microbes that produce the antibiotic or in their natural competitors [7, 8]. The high plasticity of bacterial genomes, enabled by a large repertoire of mobile genetic elements, and the availability of a large pool of ancient antibiotic resistance determinants hence set the stage for the rapid proliferation of antibiotic resistance, giving rise to multi-resistant clinical strains just a few years after the commercial introduction of antibiotics [7].

Synthetic chemotherapeutic agents predate antibiotics in the clinical setting, and continue to be used synergistically with antibiotics to treat microbial infections [9]. Following the initial discovery and clinical use of arsphenamine in 1907 [10], interest in chemotherapeutic agents quickly took off after the development of sulfa drugs in the 1930’s [11]. The discovery of trimethoprim (a di-aminopyrimidine) was received with interest because, like sulfonamides, trimethoprim targets the bacterial synthesis of tetrahydrofolic acid, which is a necessary cofactor in the synthesis of thymine and purines [12]. Sulfonamides are structural analogs of para-aminobenzoic acid (PABA) and inhibit the synthesis of di-hydropteroate by competing with PABA for binding to the di-hydropteroate synthase (DHPS) enzyme, resulting in sulfonamide-bound di-hydropterin [13]. Trimethoprim is a structural analog of di-hydrofolic acid, derived from di-hydro-pteroate. It acts by competitively binding to the di-hydrofolate reductase (DHFR) enzyme, and hence inhibiting the production of tetrahydrofolic acid [13, 14]. Hence, the synergistic use of trimethoprim and sulfonamides was expected to have a potent bactericidal effect by producing a serial blockade on the tetrahydrofolic acid pathway [12, 15].

Unlike antibiotics, chemotherapeutic agents are not produced by natural organisms, yet resistance to these novel drugs arose quickly after their mass-production and it is today pervasive among clinical isolates [7]. In the case of sulfonamides and trimethoprim, which are usually administered in tandem, resistance via chromosomal mutations to both chemotherapeutics was reported soon after their clinical introduction [13]. Chromosomal resistance to sulfonamides can occur through mutations yielding increased production of PABA [16] or, more commonly, via mutations to the chromosomal *folP* gene (encoding DHPS), which decrease the affinity of DHPS for sulfonamide without detriment to PABA binding [13, 17]. Such mutations have been reported in multiple bacterial groups and target different conserved regions of DHPS [13]. Similarly, chromosomal resistance to trimethoprim may arise via mutations that increase transcription of the *folA* gene (encoding DHFR) [18], or through mutations that decrease the affinity of DHFR for trimethoprim [13]. The vast majority of resistant clinical isolates to both sulfonamides and trimethoprim, however, are not due to chromosomal mutations, but to the acquisition of resistance determinants on mobile genetic elements [13]. Parallel to their systematic combined use in both clinical and agricultural settings, genes conferring resistance to sulfonamides and trimethoprim are frequently found together on mobile elements, such as class 1 integrons [19] or conjugative plasmids [13, 20]. The mobile genes conferring resistance to sulfonamide are homologs of the chromosomally-encoded *folP* gene and are collectively known as *sul* genes (for *sul*fonamide resistance). Mobile genes conferring resistance to trimethroprim are either homologs or functional analogs of the chromosomally-encoded *folA* gene and are collectively known as *dfr* genes (for *d*i-hydro*f*olate *r*eductase) [17].

In spite of their frequent co-occurrence on mobile genetic elements, there are significant differences between the mobile genes conferring resistance to sulfonamides (*sul* genes) and trimethoprim (*dfr* genes). To date, only three *sul* gene classes have been described in clinical isolates [21], whereas more than 30 different *dfr* genes have been reported in clinically-relevant strains [22]. Trimethoprim resistance (*dfr*) genes have been further classified into two families (*dfrA* and *dfrB*). These two families encode evolutionarily unrelated proteins of markedly different sizes. Sequence similarity indicates that *dfrA* genes are homologs of the chromosomally-encoded *folA* genes, whereas *dfrB* genes are functional analogs of unknown origin [23, 24]. Most *dfrA* genes follow a standard naming convention consisting of *dfrA* followed by a numerical value indicating their discovery rank order. However, several *dfrA* genes first identified in Gram-positive bacteria, and thought at the time to be unrelated to the Gram-negative *dfrA* genes, were originally named following an alphabetical convention (*dfrC-K*). The disparity in genetic diversity among sulfonamide and trimethoprim mobile resistance determinants is suggestive of different evolutionary processes leading to the onset and spread of resistance to these two chemotherapeutic agents [13]. It was suggested that resistance to sulfonamide had arisen in a few isolated organisms and rapidly spread upon the introduction of sulfa drugs, whereas trimethoprim resistance had independently evolved, and had been subsequently mobilized, multiple times [13].

Recently, we examined the origins of *sul* genes through comparative genomics, phylogenetic analysis and *in vitro* assays [25]. We identified a well-defined mutational signature in *sul*-encoded proteins with respect to chromosomally encoded *folP* genes, and we used this conserved motif to map the origins of *sul* genes in bacterial chromosomes. Our work revealed that the three groups of *sul* genes identified in clinical isolates originated in the *Leptospiraceae* and were transferred to the *Rhodobiaceae* more than 500 million years ago. These two ancient resistant determinants were later independently mobilized, and rapidly disseminated following the commercial introduction of sulfa drugs. By tracing the phylogenetic lineage of *sul* genes and demonstrating that these two bacterial families were resistant to sulfonamides long before their discovery and clinical use, our work indicated that resistance to novel drugs could very well preexist, and be ready for mobilization, within the vast bacterial pangenome. Here we apply similar methods to examine the phylogenetic relationships among *dfrA* and chromosomally-encoded *folA* genes. Our aim is to shed light on the evolutionary processes giving rise to mobile trimethoprim resistance genes. Our work illustrates significant similarities and differences in the processes leading to the emergence and spread of trimethoprim and sulfonamide resistance determinants, reveals previously unreported clusters of *dfrA* genes, and suggests that systematic analyses of the bacterial pangenome may of use in the design of novel antibacterials.

## 6. Methods

### 6.1 Sequence data collection

To identify homologs of DfrA proteins, we first compiled a panel of Dfr proteins reported in the literature. Dfr proteins belong to two distinct families of unrelated sequences (DfrA and DfrB; Figure S1). We mapped these sequences to PFAM models of DfrA (PF00186) and DfrB (PF06442) using HMMER (hmmscan), and we discarded sequences mapping to the DfrB family, retaining only DfrA proteins for analysis (Table S1; Data 1). We further excluded redundant DfrA sequences (identity >90%) using T-COFFEE seq_reformat command [26], and used the resulting non-redundant panel to identify DfrA homologs in protein records associated with NCBI GenBank/RefSeq sequences corresponding to mobile genetic elements. These were defined as sequences containing the keywords “plasmid”, “integron” or “transposon” in their title, belonging to complete genome records [27, 28]. BLASTP hits matching stringent e-value (<1e^−20^) and query coverage (>75%) thresholds were added to the panel if non-redundant (identity <90% with respect to existing panel members), and their classification as mobile elements was validated by assessing that they contained at least one gene coding for an integrase, transposase or plasmid replication protein, as determined by HMMER (hmmscan, e-value<1e^−05^) with reference PFAM models (Table S2) [29–33]. To detect putative chromosomally-encoded *folA* genes associated with mobile *dfrA* genes, we used the sequences in the extended non-redundant panel of DfrA homologs as queries for TBLASTN searches against NCBI GenBank complete genomes with stringent e-value (<1e^−40^) and query coverage (>75%) settings. Hits with nearby genes annotated as resistance determinants, transposases or integrases were considered to be chromosomally-integrated mobile DfrA homologs and not considered for further analysis. For each mobile DfrA homolog in the panel, the first, if any, TBLASTN hit satisfying these thresholds was considered, for the purposes of this study, to be a proxy for the closest putative chromosomally-encoded FolA protein. The choice of representative DfrA sequences did not alter the TBLASTN results. To complete the panel of sequences used to reconstruct the evolutionary history of DfrA/FolA sequences, we used the non-redundant panel of mobile DfrA sequences to identify via BLASTP (e-value<1e^−20^, coverage>75%) FolA proteins encoded by the chromosomes of NCBI RefSeq representative species for all bacterial orders, and for each bacterial family in the Proteobacteria. In addition, the closest archaeal homologs of bacterial FolA sequences were determined by searching with BLASTP the NCBI protein database, restricted to Archaea (taxid:2157), with the *Escherichia coli* FolA protein. A member of each family from the order (Halobacteriales) of the identified best archaeal hit of *E. coli* FolA was sampled to populate the outgroup.

### 6.2 Phylogenetic analysis

For phylogenetic inference, we performed a T-COFFEE multiple sequence alignment of protein sequences for the complete panel of DfrA and FolA homologs, combining three CLUSTALW profile alignments with different (5, 10, 25) gap opening penalties and leveraging *E. coli* FolA crystal structure (P0ABQ5) to adjust gap penalties [34]. The resulting alignment was processed with Gblocks (Allowed Gap Positions: With Half, Minimum Number Of Sequences For A Conserved Position: 86, Minimum Number Of Sequences For A Flanking Position: 95, Maximum Number Of Contiguous Nonconserved Positions: 5, Minimum Length Of A Block: 4) [35]. Bayesian phylogenetic inference on the trimmed multiple sequence alignment was carried out with MrBayes version 3.2.6 [36]. Four Metropolis-Coupled Markov Chain Monte Carlo simulations with four independent chains were run for 20,000,000 generations, using a mixed four-category gamma distributed rate plus proportion of invariable sites model [invgamma] and a JTT (Jones-Taylor-Thornton) amino acid substitution model [37]. Chains were sampled every 100 iterations and stationarity was analyzed with Tracer [38] by monitoring the Estimated Sample Size (ESS). To determine burnin, chain results were summarized with MrBayes imposing the restriction that ESS be above 200 and that the Potential Scale Reduction Factor (PSRF) be within 0.005 of 1. Based on summarization results, the burn-in was set at 20% of iterations. A consensus tree was generated with the half-compat option and visualized using the GGTREE R library [39]. Ancestral state reconstruction of a single binary trait (mobile/chromosomal) was performed with BayesTraits version 3.0.2 [40]. Known states at tree tips were labeled, and ancestral states were reconstructed using the Multistate and Maximum Likelihood (ML) settings.

### 6.3 DNA Techniques and In vitro Trimethoprim Susceptibility Assay

With the exception of the *Ralstonia solanacearum* GMI1000 (Marc Valls; Center for Research in Agricultural Genomics) and *E. coli* K-12 (CGSC5073) *folA* genes, which were amplified from genomic DNA, *dfrA* and *folA* homologs were adapted to *E. coli* codon usage, synthesized (ATG:biosynthetics GmbH) and then subcloned into a dephosphorylated pUA1108 vector [41] using NdeI and BamHI double digest (New England Biolabs) when possible. Genes with internal restriction sites for any of these two enzymes were subcloned into the same vector using the HIFI DNA Assembly kit (New England Biolabs) following the manufacturer’s protocol. Oligonucleotides used in this work are listed in Table S3. All constructs were validated by sequencing (Macrogen) prior to their transformation into *E. coli* K-12 (CGSC5073). The minimal inhibitory concentration (MIC) for trimethoprim (Sigma-Aldrich) for strains of *E. coli* K-12 (CGSC5073) carrying different versions of pUA1108 encoding *folA* or *dfrA* homologs was determined following the Clinical and Laboratory Standards Institute (CLSI) guidelines using microdilution tests in Mueller-Hinton broth (Merck) [42]. All MIC assays were performed in triplicate. Colonies were grown on Luria-Bertani (LB) agar for 18 h and then suspended in sterile 0.9% NaCl solution to a McFarland 0.5 turbidity level. Suspensions were then diluted at 10^−2^ in Mueller-Hinton (MH) broth, and 50 μl (5×10^4^ cells) were inoculated onto microtiter plates that contained 50 μl of MH broth supplemented with 1024–0.250 mg/L of trimethoprim. To determine growth, absorbance at 550 nm was measured after 24 h incubation at 37°C. The *dfrA1* gene was used as positive control [43] and the *E. coli folA* gene as a negative control [44].

### 6.4 Sequence analysis

To assess whether the identified chromosomal gene associated with a mobile *dfrA* gene is the canonical *folA* gene for the genus, and not the product of a subsequent recombination of the mobile *dfrA* gene into the chromosome, we computed the pair-wise amino acid identity among the products of all chromosomal *folA* homologs and then compared this distribution with the pair-wise amino acid identity of the putative origin versus the chromosomal *folA* homologs. We used a onesided Mann-Whitney U test to determine if the two distributions were significantly different. To analyze the %GC content relationship between *sul/dfrA* genes and their host chromosomes, we used pre-compiled panels of sequences for non-redundant Sul [25] and DfrA homologs to search protein records associated with NCBI GenBank/RefSeq sequences of mobile genetic elements. The %GC content of the corresponding *sul* and *dfrA* genes, as well as the overall %GC content in both the mobile genetic element and the chromosome of the species harboring it, were computed with custom Python scripts. To analyze whether mobile *dfrA* genes with %GC content close to their hosts’ genomes are more similar to the hosts’ *folA* genes than expected if *dfrA*-host associations were arbitrary, we performed a permutation test comparing the mean pairwise alignment distance between DfrA genes and host-encoded FolA genes. We randomly permuted DfrA-host assignments 1,000 times and computed the corresponding p-value as the rank of the non-permuted mean pairwise alignment distance. The input files (Data 2) and scripts (Data 3) used for data collection and analysis are available on public repositories.

## 7. Results and discussion

### 7.1 A large fraction of reported *dfrA* genes share a common evolutionary origin

To explore the phylogenetic relationship of trimethoprim resistance determinants with their chromosomally-encoded *folA* counterparts, we used a non-redundant panel (<90% pairwise identity) of protein sequences encoded by reported *dfr* genes (Table S1) to detect putative DHFR homologs in sequenced mobile elements. We mapped all these reference sequences to the PFAM models for DfrA and DfrB (Table S4) [45]. We discarded sequences associated with the *dfrB* gene family, and retained for analysis non-redundant sequences mapping to the clades defined by *dfrA* genes reported in the Proteobacteria and by *dfrDEFGK* genes associated with Gram-positive bacteria. For convenience, and in accordance with recent reports on *dfr* nomenclature [46], we hereinafter designate these two groups (*dfrA* and *dfrDEFGK*) with the umbrella term *dfrA*. These reference mobile DfrA homologs were then used to detect putative chromosomally-encoded homologs in complete bacterial genomes using TBLASTN. These sequences were combined with FolA homologs sampled from representative genomes of all bacterial orders with complete genome assemblies, and of each bacterial family within the Proteobacteria (Table S5; Data 4). The resulting multiple sequence alignment was used to perform Bayesian phylogenetic analysis of FolA/DfrA sequences.

The phylogenetic tree shown in Figure 1 showcases the genetic diversity of *dfrA* genes, which encompass sequences with pairwise sequence identities ranging from 99% to 20% (Table S6). The resulting phylogeny also reveals that the vast majority (70.7%) of known DfrA sequences map to two well-supported (>0.8 posterior probability), distinct clades that likely arose from two different ancestors. The first clade (*Clade 1*), typified by the DfrA1 and DfrA12 proteins [47], includes 22 sequences encoded by previously reported *dfrA* genes with an average amino acid identity of 51.19% ±SD 17.63%, divided into two subgroups (containing 17 and 5 known *dfrA* genes, respectively) and associated with Gammaproteobacteria pathogens. This clade includes also a basal set of taxa composed of the *Clostridioides difficile dfrF* gene and two newly identified mobile *dfrA* homologs also from Firmicutes isolates. The second clade (*Clade 2*), exemplified by DfrA18 [48], comprises a group of six highly diverged (34.37% ±SD 10.15% average amino acid identity) DfrA sequences from Gammaproteobacteria isolates.

**Figure 1.**
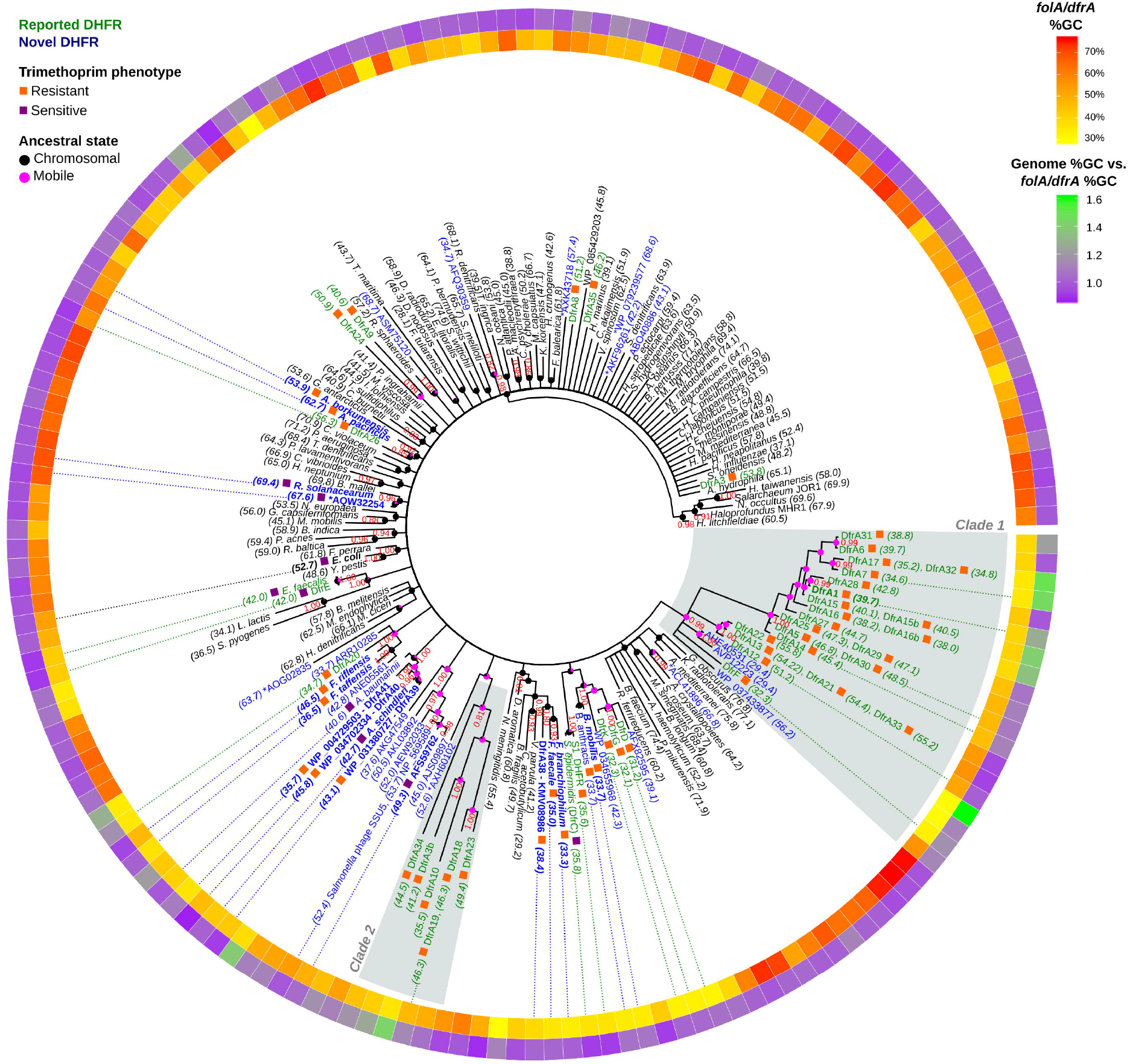
Consensus tree of DHFR protein sequences. Branch support values are provided as Bayesian posterior probabilities estimated after four independent runs of 20,000,000 generations. Support values are only shown for branches with posterior probability values higher than 0.8. For chromosomal DHFR, the species name is displayed. Mobile DHFRs are denoted by their established *dfrA* name or by their NCBI GenBank accession numbers. Reported *dfrA* genes deemed redundant (>90% identity) are listed next to the corresponding non-redundant taxon included in the analysis. Next to each tip label, colored boxes designate (orange) trimethoprim resistant and (purple) sensitive DHFR. Numbers between brackets indicate the %GC content of the sequence for the gene encoding the DHFR. Tip label coloring denotes: (green) previously reported and (blue) novel DHFRs. Bold label text indicates that resistance has been experimentally assessed in this work. DHFR variants marked with an asterisk (*) are encoded in megaplasmids (> 400 kbp). The internal ring shows the %GC of the gene encoding the DHFR in yellow-red color scale, while the external ring displays the ratio between the %GC content of the genome harboring the DHFR gene and the %GC content of the gene. Dotted lines from the inner ring to tip labels denote genes discussed in the text. Reconstructed mobile/chromosomal states are displayed on ancestral nodes as pink/black pie charts.

Analysis of the *dfrA* gene sequences in these two clades reveals an unexpected degree of heterogeneity in %GC content. In the first clade, several *dfrA* homologs, including the *C. difficile dfrF* gene, show relatively low %GC content (Figure 1 inner ring), matching the Firmicutes species they were reported on (Figure 1 outer ring; Table S7). Similarly, *dfrA* genes in the *dfrA12* group show a %GC content (53.28 ±SD 1.80) that is well in line with that of the *Enterobacteriaceae* isolates harboring them. Conversely, the largest group in this clade, encompassing *dfrA1, dfrA7* and *dfrA14*, shows a mean %GC content of 41.03 ±SD 3.99, which is substantially lower than the average %GC content of the *Enterobacteriaceae* harboring these mobile elements. The same holds true for the second clade (*dfrA18*), which also shows lower %GC content (43.88 ±SD 4.91) than expected for the *Enterobacteriaceae*. To ascertain whether this pattern of %GC heterogeneity extended to other previously reported and putative *dfrA* genes, we examined the %GC content of *dfrA* (935) and *sul* (408) gene homologs identified in this analysis with respect to the genome %GC content of the host species harboring these mobile resistance genes.

The results shown in Figure 2 (top panel) and Table S8 (Data 5) reveal that *dfrA* genes tend to align with host genome %GC content (Pearson ρ=0.56), whereas *sul* genes display a two-tiered distribution of %GC content that is essentially independent of host genome %GC (Pearson ρ=0.14). Available *dfrA* and *sul* sequences are dominated by variants of a known *dfrA* and *sul* genes that have been isolated predominantly in a select group of bacterial hosts (Figure 2; bottom panel). To correct for this skew, we filtered *dfrA* sequences based on the amino acid identity (<90%) of their encoded proteins. This filtering resulted in a significantly smaller number of non-redundant representative *dfrA* (63) and *sul* (4) genes (Table S9). The four representative *sul* genes correspond to one exemplar of the *sul1* and *sul2* families, and two exemplars of the *sul3* family. Among representative *dfrA* genes, fourteen map to the first clade (*Clade 1*) of Figure 1 and four to the second clade (*Clade 2*). The correlation of *dfrA* genes with host genome %GC increases significantly (Pearson ρ=0.78) when considering only non-redundant representative *dfrA* sequences. The fact that the %GC content of representative *dfrA* sequences aligns well with their host genome %GC could suggest that %GC content in *dfrA* genes has been ameliorated to match the host’s. Alternatively, it could indicate that the mobile *dfrA* gene originated via mobilization of a chromosomal *folA* gene from a bacterium in the same clade as the current host. The later scenario posits that, besides %GC content similarity, representative *dfrA* genes should also encode proteins with significant sequence similarity to their hosts’ FolA protein. We performed a permutation test to analyze whether representative *dfrA* gene products show significant similarity with their hosts’ FolA protein (Table S10). Our results indicate that this is the case (p<0.001), suggesting that most mobile *dfrA* genes are still associated with species from the same clade they originated in.

**Figure 2.**
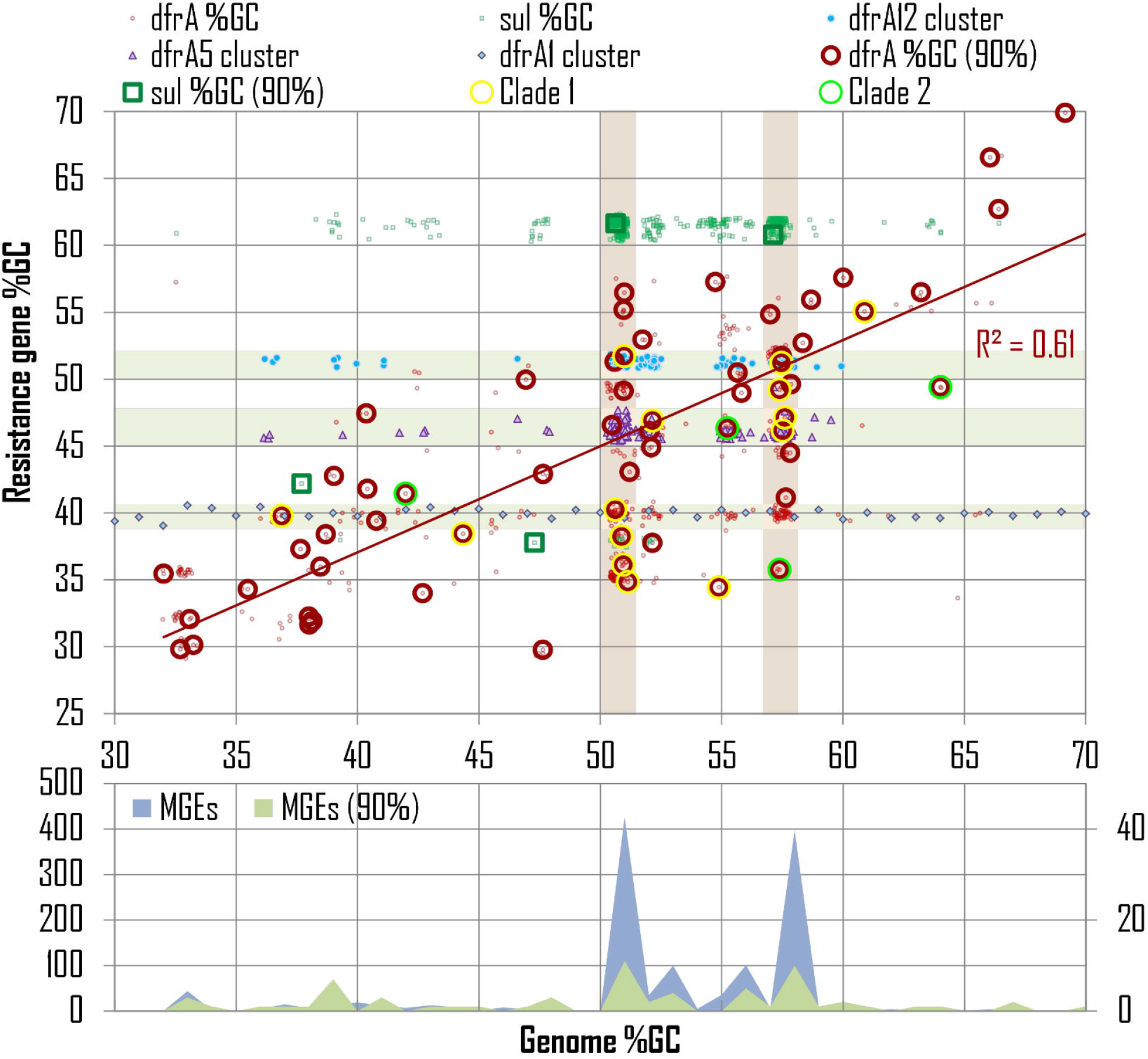
Correlation between the %GC content of mobile *dfrA* (red circles) and *sul* (green squares) genes and that of their host genome. Large open circles/squares denote representatives of clusters of redundant sequences (identity >90%), and *dfrA* genes from **Figure 1** *Clade 1* and *Clade 2* are marked with an additional corona. A 0.75% jitter to both x- and y-axis values has been applied for visualization purposes. The red line shows the linear regression for representative *dfrA* gene values. The Pearson R^2^ coefficient is superimposed. Vertical background bars in the top panel designate DfrA sequences harbored by MGEs identified in *E. coli* and *K. pneumoniae* isolates, which are heavily overrepresented in the dataset. Sequences from clusters with more than 100 sequences (represented by *dfrA12, dfrA5* and *dfrA1*) are shown with specific markers, and highlighted by horizontal background bars. The number of mobile genetic elements (MGEs) identified as harboring *dfrA* genes, before and after filtering DfrA sequence identity (>90%), is shown in the bottom panel.

The filtering of *dfrA* sequences based on amino acid identity generates three large clusters (represented by *dfrA12, dfrA5* and *dfrA1*, and belonging to *Clade 1* from Figure 1) containing more than 100 genes with identity larger than 90%. The *dfrA* genes in these clusters show a distribution of %GC content that is essentially independent of the host genome %GC, as in the case of *sul* genes (Figure 2; top panel), and their products show no significant sequence similarity with the hosts’ FolA (permutation test p>0.1). This indicates that the *dfrA* genes in these large clusters have spread across distantly related bacterial clades, primarily through their association with *sul*-containing integron-based transposable elements that are widely disseminated among clinically relevant bacteria [49, 50]. This analysis also brings to the fore the presence of multiple *dfrA* cluster representatives, with widely divergent %GC, on narrow bands of host genome %GC content. These bands correspond to *E. coli* (50.7% GC) and *Klebsiella pneumoniae* (57.4% GC) isolates, which are heavily oversampled in the dataset (Figure 2, bottom panel). The marked divergence in %GC content (*E. coli:* 9.55% ±SD 6.15%, *K. pneumoniae:* 6.93% ±SD 4.63%) and amino acid sequence identity (*E. coli:* 38.57% ±SD 15.24%, *K. pneumoniae:* 38.39% ±SD 14.02%; Table S11; Figure S2) among these representative *dfrA* genes suggests that they originated via mobilization from a diverse set of chromosomal backgrounds.

The fact that the %GC of *dfrA* genes aligns with their host genome’s %GC and that DfrA proteins display higher sequence identity when aligned to their host genome FolA proteins than to other FolA proteins strongly supports the notion that *dfrA* genes have been mobilized multiple times within different bacterial clades [13]. In a few instances, typified by the large *dfrA* clusters illustrated in Figure 2, *dfrA* genes have been captured by highly efficient mobile elements and dispersed widely across unrelated groups of bacteria [49]. These mobile elements often harbor *sul* genes, which also display a host-independent %GC distribution. Many of the *dfrA* genes identified here are associated with clinical isolates. The divergent %GC content and amino acid identity of these *dfrA* genes indicates that pathogenic bacteria have obtained *dfrA* genes on multiple occasions and from different sources, highlighting the ability of mobilized resistance determinants to reach clinically-relevant pathogens [17, 51].

### 7.2 Novel trimethoprim resistance determinants of *Acinetobacter* clinical isolates identified through phylogenetic methods

The phylogenetic tree in Figure 1 includes reported DfrA proteins and their putative homologs, as well as FolA proteins identified via TBLASTN as putative DfrA homologs or sampled uniformly across bacterial clades. The inferred phylogeny also reveals several groups of previously unreported mobile DHFR homologs that form well-supported clades in association with chromosomal FolA proteins. These FolA proteins could hence constitute the chromosomal origins of the associated mobile DHFR homologs, and provide insights into the emergence and dissemination of trimethoprim resistance genes. To determine whether these mobile DHFR homologs did confer resistance to trimethoprim, we cloned a subset of the genes encoding them and we performed broth microdilution assays to determine the minimal inhibitory concentration (MIC) of trimethoprim. The results, shown in Table 1, reveal that most of the mobile DHFR homologs identified here do confer significant resistance to trimethoprim. The sole exception is AQW32254. Close inspection revealed that this DHFR homolog is encoded by a megaplasmid (1.2 Mb) from a *Ralstonia* isolate, and that this is the only DHFR homolog present in its complete genome. We hence determined that this DHFR homolog was a bona fide FolA protein and not a mobile DHFR homolog, and we did not consider as putative *dfrA* genes all other DHFR homologs identified in megaplasmids (> 400 kbp).

**Table 1.** Minimum inhibitory concentrations (MICs) of trimethoprim for wild-type *Escherichia coli* K-12 (CGSC5073) and derivatives carrying different versions of *dfr/folA* or the control empty vector. Values are representative of four independent replicates.

| Strain | Mobile / Chromosomal | Nucleotide accession | Cloned Protein ID | Trimethoprim (mg/L) |
| --- | --- | --- | --- | --- |
| *E. coli* CGSC5073 | - | - | - | 0.25 |
| *E. coli* pUA1108 | - | - | - | 0.25 |
| *E. coli* pUA1108::*foIA E. coli* | C | NC_000913 | WP_000624375 | 4 |
| *E. coli* pUA1108::*dfrA1* | M | NC_002525 | WP_000777554 | >512 |
| *E. coli* pUA1108::*foIA Flavobacterium branchiophilum* | C | NC_016001 | WP_014083133 | 256 |
| *E. coli* pUA1108::*foIA Flavobacterium faecale* | C | NZ_CP020918 | WP_108740183 | >512 |
| *E. coli* pUA1108::*dfrA38 Acinetobacter baumannii* | M | CP021344 | KMV08986 | 256 |
| *E. coli* pUA1108::*foIA Acinetobacter schindleri* | C | NZ_CP025618 | WP_004813248 | 0.25 |
| *E. coli* pUA1108::*dfrA39 Acinetobacter baumannii* | M | NZ_CP021785 | WP_031380727 | 512 |
| *E. coli* pUA1108::*dfrA40 Acinetobacter baumannii* | M | NZ_JEVW01000010 | WP_034702334 | 128 |
| *E. coli* pUA1108::*dfrA41 Acinetobacter defluvii* | M | NZ_CP029396 | WP_004729503 | >512 |
| *E. coli* pUA1108::*foIA Fluviicola taffensis* | C | NC_015321 | WP_013685591 | >512 |
| *E. coli* pUA1108::*foIA Candidatus Fluviicola riflensis* | C | CP022585 | ASS49886 | >512 |
| *E. coli* pUA1108::*foIA Alcanivorax pacificus* | C | NZ_CP004387 | WP_008736147 | 32 |
| *E. coli* pUA1108::*foIA Alcanivorax borkumensis* | C | AM286690 | CAL17791 | 16 |
| *E. coli* pUA1108::*foIA Bacillus mobilis* | C | NZ_CP031443 | WP_000637217 | >512 |
| *E. coli* pUA1108::*foIA Ralstonia solanacearum* | C | NC_003295 | WP_011000898 | 0.5 |
| *E. coli* pUA1108::*foIA* blood disease bacterium A2-HR MARDI | M | CP019912 | AQW32254 | 1 |
| *E. coli* pUA1108::*foIA E. coli* O104:H4 | M | CP003298 | AFS59762 | 2 |

Two remaining clades of novel mobile DHFR homologs from clinically-relevant bacteria associated with chromosomal FolA proteins were shown to confer resistance to trimethoprim on *E. coli* (Table 1). To investigate whether the sequence determinants conferring resistance had originated in the associated chromosomal background, we cloned the most closely related chromosomal *folA* gene as well as gene encoding an additional DHFR homolog from the same genus and performed broth microdilution assays to determine the MIC of trimethoprim. We also performed ancestral state reconstruction of the molecule encoding the DHFR homologs (chromosomal/mobile trait) (Table S12). The combined results of Table 1 and Figure 1 reveal different patterns of trimethoprim resistance acquisition. KMV08986 is a DHFR homolog harbored by a conjugative plasmid from an *Acinetobacter baumannii* clinical isolate. Its most closely related chromosomally-encoded DHFR homolog is the FolA protein of *Flavobacterium branchiophilum*, which confers resistance to trimethoprim (Table 1).

To ascertain whether this chromosomally-encoded DHFR homolog was encoded by a bona fide *folA* gene, instead of a mobile *dfrA* gene that integrated into the chromosome, we compared the genus-wide distribution of pairwise alignment distances between FolA proteins to the pairwise distance of the identified homolog versus all other FolA proteins in the genus. The *F. branchiophilum* FolA sequence is significantly different from other *Flavobacterium* FolA sequences (Mann Whitney U p<0.05; Table S13), raising the possibility that this chromosomal gene could be in fact a recombined mobile *dfrA* gene. However, phylogenetic analysis with a broader representation of *Flavobacterium* sequences (Figure S3) confirms the well-supported branching of *F. branchiophilum* FolA with other *Flavobacterium* species FolA proteins, and comparative genomics analysis reveals that the genetic neighborhood of the chromosomal *folA* gene is conserved in the *Flavobacterium* genus (Figure S4). Furthermore, the FolA protein of a prototypical genus member, *Flavobacterium faecale*, also confers resistance to trimethoprim on *E. coli* (Table 1). These results indicate that the FolA protein was likely resistant to trimethoprim in the ancestor of extant *Flavobacterium* species, which diverged more than 50 million years ago [52]. The branching of KMV08986 in the reconstructed phylogeny and the associated ancestral state reconstruction indicates that this mobile DHFR homolog likely originated via mobilization of a chromosomal *folA* gene within the Bacteroidetes phylum. The encoded FolA protein was likely resistant to trimethoprim, but the exact donor species remains to be elucidated.

In contrast to *Flavobacterium* proteins, the *Acinetobacter schindleri* FolA protein does not confer resistance to trimethoprim, in agreement with previous reports of *A. schindleri* susceptibility to trimethoprim [53], and with the well-established susceptibility of *A. baumannii* FolA to trimethoprim [54, 55]. The *A. schindleri* FolA protein is closely related to three mobile DHFR homologs conferring resistance to trimethoprim and harbored by *A. baumannii* (WP_031380727, WP_034702334) and *Acinetobacter defluvii* (WP_004729503) clinical and environmental isolates. These mobile DHFR homologs branch within a well-supported clade of chromosomal *Acinetobacter* FolA proteins, as supported by ancestral state reconstruction (Figure 1; Table S12). The trimethoprim susceptibility of *Acinetobacter* chromosomal *folA* genes and the phylogenetic placement of these DHFR homologs hence indicates that the observed resistance to trimethoprim was acquired immediately prior to or after mobilization from an *Acinetobacter* chromosomal background. This is supported by the observation that these mobile DHFR homologs confer different levels of resistance to trimethoprim (Table 1), and that the largest MIC correlates with the location of the DHFR homolog on a plasmid harboring multiple antibiotic resistance determinants (Figure S5). This suggests that these DHFR homologs have acquired mutations conferring heightened resistance to trimethoprim in parallel to their broader dissemination on multi-resistant mobile elements. Based on their validated trimethoprim resistance phenotype and their level of sequence identity versus previously reported DfrA proteins (<95%; Table S14) [13], we propose to designate these *Acinetobacter* DHFR homologs as DfrA38 (KMV08986), DfrA39 (WP_031380727), DfrA40 (WP_034702334) and DfrA41 (WP_004729503).

Here we report the identification of trimethoprim susceptible chromosomal *folA* genes that are closely related to mobile *dfrA* genes, as well as the discovery of chromosomally-encoded *folA* genes conferring resistance to trimethoprim. This indicates that, in contrast to sulfonamides [25], trimethoprim resistance mutations with small or negligible fitness cost must occur frequently enough in natural environments. These *folA* variants can then be selected for and mobilized upon exposure to trimethoprim. It is well-documented that resistance to trimethoprim, mediated by mutations on the chromosomal *folA* gene, develops very rapidly and in a fairly structured way [56–58], whereas resistance to sulfonamides takes much longer to evolve in a laboratory setting. Moreover, sulfonamide resistant mutants typically show significantly reduced affinity to PABA. This results in a net fitness cost in the absence of sulfonamide that is only palliated by the emergence of subsequent compensatory mutations [59, 60]. Beyond structural constraints on the respective binding pockets, a crucial difference between both chemotherapeutic agents lies in their respective targets. While trimethoprim directly inhibits DHFR, sulfonamides compete with PABA for access to DHPS, yielding a non-productive sulfonamide-bound di-hydropterin. For sulfonamides, therefore, it is the PABA-to-sulfonamide ratio that limits the production of di-hydropteroate from a limited pool of pteridine di-phosphate, and this cannot be altered via overexpression of DHPS [61]. Conversely, trimethoprim overexpression can provide partial resistance to trimethoprim, and mutations enhancing DHFR expression have been reported to be the first to appear in directed evolution experiments [58]. The ability to obtain partial resistance through overexpression may hence provide a stepping stone for the gradual accumulation and refinement of mutations conferring substantial resistance with little fitness cost, and hence facilitate the development of trimethoprim resistance [57, 58].

### 7.3 Trimethoprim resistance in chromosomally-encoded *folA* genes

Besides uncovering novel *dfrA* genes, the phylogenetic analysis in Figure 1 also identifies several chromosomal *folA* genes associated with previously reported *dfrA* genes. Two of these chromosomal *folA* genes have already been reported in the literature as putative origins of *dfrA* genes, and their identification here provides some degree of validation for the phylogenetic approach implemented in this work. The putative chromosomal origin for *Staphylococcus aureus Tn4003* S1-DHFR has been identified as the chromosomally-encoded *dfrC* gene (*Staphylococcus epidermidis*) and reported to be susceptible to trimethoprim [62]. The *Enterococcus faecalis dfrE* gene, identical to the chromosomally-encoded *folA* gene of *E. faecalis*, was reported to confer moderate resistance to trimethoprim in *E. coli*, but only when cloned in a multicopy plasmid, which could easily result in overexpression-mediated resistance [61, 63].

To ascertain whether the chromosomal *folA* genes found here to be associated with other known *dfrA* genes (*dfrA20, dfrA26* and the *dfrDGK* cluster) confer resistance to trimethoprim, we performed broth microdilution assays to determine the MIC of trimethoprim on these chromosomally-encoded FolA proteins and on another FolA protein from the same genus. In all cases, both related FolA proteins confer resistance to trimethoprim (Table 1). The most closely associated chromosomal *folA* genes are not significantly different from other *folA* genes in their respective genera (Mann Whitney U p>0.05; Table S13), as reflected also by substantial conservation of the *folA* genomic neighborhood (Figure S4). Together, these data indicate that resistance to trimethoprim was present on the ancestor of these genera. The *dfrA26* gene was identified on a *K. pneumoniae* clinical isolate and its most closely associated chromosomal *folA* gene is a member of the *Alcanivorax* genus. The branching pattern of *dfrA26* within this clade and ancestral state reconstruction results (Figure 1; Table S12) suggest that it arose via mobilization of a chromosomal *folA* gene from the *Alcalinivorax* genus. The *dfrDGK* genes have been reported, respectively, in *E. faecalis, Enterococcus faecium* and *S. aureus*, and ancestral state reconstruction results indicate that these mobile *dfrA* genes originated through mobilization of a member of closely-related the *Bacillus* genus, members of which have been reported to be naturally resistant to trimethoprim [64]. In both cases, therefore, the phylogenetic evidence and the similarity in %GC content among chromosomal and mobile genes (Figure 1, Table S15) point towards a mobilization event that has to date remained circumscribed to related genera. Conversely, the *dfrA20* gene was identified on *Pasteurella multocida* isolate, yet the chromosomal *folA* gene most closely associated to it is encoded by *Fluviicola taffensis*, a Bacteroidetes, hence suggesting a much more distant mobilization event (Figure 1, Table S15). In all three cases, however, we find evidence that preexisting resistant *folA* genes can be readily mobilized from both close (e.g. *dfrDGK*) or distant (e.g. *dfrA20*) species.

The resistance to trimethoprim reported here for the chromosomal *folA* genes of two different genera of Bacteroidetes, two distinct *Alcanivorax* species and a *Bacillus* strain underscores the deep ancestry of chromosomal mutations yielding resistance to trimethoprim. The *folA* genes of *Flavobacterium* and *Fluviicola* were shown here to confer resistance to trimethoprim. These two genera are thought to have diverged more than 500 milion years and define major lineages within the Flavobacteriales, suggesting that resistance to trimethoprim emerged in an ancestor of this bacterial order. It is worth noting that several of the chromosomal *folA* genes shown here to be associated with mobile DHFR homologs (*Alcanivorax, Flavobacterium* and *Fluviicola*) appear to be resistant at the genus level and correspond to genera of aquatic bacteria. This parallels our recent identification of soil and subterranean water bacteria as the likely originators of clinical sulfonamide resistance genes [25], and suggests that the intensive use of trimethoprim/sulfamethoxazole in agriculture, aquaculture and animal husbandry in the last fifty years may have acted as a trigger for the selection and mobilization of preexisting *folA* and *folP* genes conferring resistance to trimethoprim and sulfonamides. Conversely, trimethoprim susceptible chromosomal *folA* genes found here to be associated with *dfrA* genes belong to clinically-relevant genera (*Staphylococcus* and *Acinetobacter*) that may have been under more direct trimethoprim pressure. This suggests that among relatively isolated bacterial populations frequent exposure to high levels of trimethoprim may trigger the mobilization of spontaneous *folA* mutants, whereas longer term exposure to sub-lethal doses of trimethoprim in ecological rich habitats might instead rely predominantly on the mobilization of naturally resistant *folA* genes (Figure 3).

**Figure 3.**
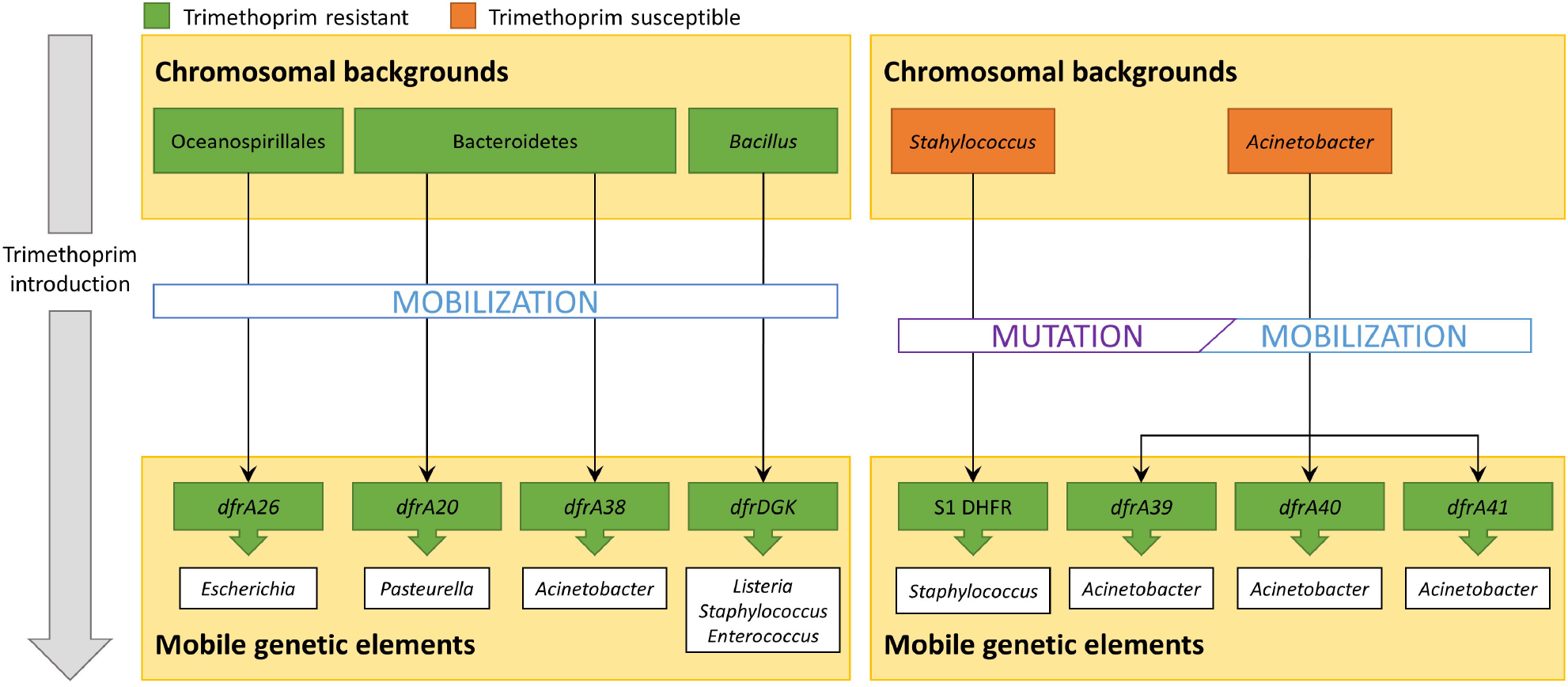
Schematic representation of the two proposed evolutionary processes, based on the results presented in **Figure 1**, **Figure 2** and **Table 1**, leading to the dissemination of trimethroprim resistance determinants. (Left panel) Upon the introduction of trimethoprim, mobilization events involving preexisting resistant chromosomal *folA* genes can be favorably selected. (Right panel) Following the introduction of trimethoprim, mobilization events involving *folA* genes with novel mutations that confer resistance to this chemotherapeutic agent may be selected for and disseminated among closely related bacteria.

### 7.4 Phage-encoded *folA* genes do not confer resistance to trimethoprim

Our phylogenetic analysis also identifies a well-defined clade of *Enterobacteriaceae* cryptic plasmids derived from *Salmonella phage* SSU5 and encoding DHFR homologs [65–68]. Genes coding for DHFR homologs occur frequently in many bacteriophage families, often in tandem with thymidylate synthase genes [69], but their functional role has not been fully elucidated. We performed broth microdilution assays to determine the MIC of trimethoprim of *E. coli* O104:H4 DHFR (AFS59762). This phage-encoded DHFR does not confer resistance to trimethoprim (Table 1). The high sequence identity and neighborhood conservation among the DHFR enzymes encoded by these *Enterobacteriaceae* cryptic plasmids and phages (Table S16, Figure S4) therefore suggests that all these DHFR enzymes are susceptible to trimethoprim.

Bacteriophages can transfer substantial amounts of genetic material via generalized transduction, and their potential as reservoirs of antibiotic resistance determinants has gained increased attention with the advent of metagenomics [70, 71]. However, recent studies have shown that many potential resistant determinants encoded by phages do not confer resistance against their putative targets. Furthermore, only a small proportion of complete phage genomes contain putative antibiotic resistance genes [72]. Enzymes participating in the folate biosynthesis pathway, however, are relatively frequent in phage genomes. These include homologs of the *folP* gene encoding DHPS, of the *thyX* gene encoding flavin-dependent thymidylate synthase [73–75] and, predominantly, homologs of the *folA* gene encoding DHFR often found in tandem with the *thyA* gene encoding type 1 thymidylate synthase [69].

Early work on *Enterobacteria* phage T4 showed that the phage-encoded *thyA* and *folA* gene products are functional and participate also in the phage baseplate structure [76], and *thyX* has been shown to be functional in a number of phages [73–75]. It has been proposed that these genes help bacteriophages overcome shortages in the deoxynucleotide pool during replication, but their potential in conferring resistance to sulfonamides or trimethoprim remains largely unexplored. The detection here of DHFR homologs in *Enterobacteriaceae* cryptic plasmids and phages, and the subsequent assessment of their trimethoprim susceptibility, reinforces the notion that these genes have been functionally co-opted by phages principally for deoxynucleotide synthesis. Nonetheless, these genes may still confer partial trimethoprim resistance as a byproduct of *folA* overexpression, as recently reported for *Stenotrophomonas maltophilia* phage DLP4 [77].

### 7.5 Conclusions

Recent work has shown that resistance to sulfonamide, a synthetic chemotherapeutic agent, can be present in the bacterial pangenome well before their discovery. Here we have used a combination of *in silico* and *in vitro* techniques to identify novel trimethoprim resistance genes and to identify chromosomal *folA* genes that are strongly associated with novel and previously reported *dfrA* genes. We find that most of the chromosomal *folA* genes associated with mobile *dfrA* genes confer resistance to trimethoprim, but we detect cases of novel mutations being rapidly mobilized. Our work hence shows that the observations from sulfonamide resistance extend to trimethoprim, with generalized chromosomal resistance determinants predating the origin of several genera and several clusters of resistance genes disseminated broadly among clinical isolates. Moreover, this work also reveals that, unlike sulfonamides, resistance to trimethoprim is relatively easy to generate and frequently associated with species from the same clade it originated in. The identification of ancient resistance determinants for two synthetic chemotherapeutic agents strongly suggests that resistance to any novel drugs is likely to be already present in the bacterial pangenome. Systematic screening of existing natural variants could therefore provide the means to preemptively identify derivatives presenting widely-distributed natural resistance determinants and, conversely, to engineer derivatives that circumvent most, if not all, natural resistant variants.

## 8. Author statements

### 8.1 Authors and contributors

Conceptualization, JB, IE; data curation, MS-O, IE; formal analysis, MS-O, IE; funding acquisition, ML, JB; investigation, MS-O, PC, IE; methodology, ML, JB, IE; project administration, PC, ML, JB, IE; resources, PC, ML, JB; software, MS-O, IE; supervision, PC, JB, IE; visualization, MS-O, IE; manuscript preparation – original draft, MS-O, IE; manuscript preparation – review and editing, MS-O, PC, ML, JB, IE.

## 8.4 Acknowledgements

The authors wish to thank Joan Ruiz and Susana Escribano for their technical support during some of the experimental procedures, as well as Ángela Martínez-Mateos for her continued support. The authors also express their gratitude to Dr. Marc Valls for kindly providing the *R. solanacearum* GMI1000 strain.

## 10. Data bibliography

**Data 1** – The compiled list of accession numbers and references of experimentally validated *dfrA* genes is available in Table S1.

**Data 2** – Input files (JSON, TXT and FASTA) and BLAST database for Python scripts used in data collection and analysis (DOI: 10.6084/m9.figshare.12156891.v1).

**Data 3** – Release of the GitHub repository containing the Python scripts used for data collection and analysis (DOI: 10.5281/zenodo.3760352).

**Data 4** – The accession numbers of chromosomal and mobile DfrA/FolA sequences used in this work are provided in Table S5.

**Data 5** – The accession numbers of mobile element complete assemblies and their DfrA/Sul-encoded proteins used for %GC analysis are provided in Table S8.

## 12. Supplementary material

### 12.1 Supplementary Figures

**Figure S1** – Multiple sequence alignment including all reported mobile DHFR proteins. DfrB protein sequences are highlighted in yellow.

**Figure S2** – Pairwise percent amino acid identity and %GC difference between aligned representative DfrA protein sequences harbored by mobile genetic elements of *E. coli* and *K. pneumoniae*.

**Figure S3** – Unrooted Neighbor-Joining tree of DHFR protein sequences. Branch support values are provided as the total number of 1000 bootstrap pseudo-replicates in which the branching was observed. Support values are only shown for branches with at least 75% bootstrap support. The DfrA20, DfrA38, *Fluviicola* and *Flavobacterium* DHFR protein sequences are highlighted.

**Figure S4** – Schematic representation of the genetic environment of mobile DHFR genes, their putative chromosomal origin and one representative complete genome assembly for each species within the corresponding genus. Arrow boxes indicate coding regions (discontinued arrows pinpoint pseudogenes). When available, gene names or NOG identifiers are provided and color coded.

**Figure S5** – Graphical overview of *Acinetobacter defluvii* plasmid pOXA58_010030 with *dfrA41* (red arrow box) and other resistance determinants represented as blue arrow boxes. This figure was constructed using SnapGene Viewer.

### 12.2 Supplementary Tables

**Table S1** – List of *dfrA* genes reported in the literature.

**Table S2** – Proteins mapping to replicative functions of mobile genetic elements identified via HMMER search with reference PFAM domains in MGEs containing putative *dfrA* genes.

**Table S3** – List of oligonucleotides used in this work.

**Table S4** – Table S4 - HMMER mapping between the product of *dfrA* genes and PFAM models corresponding to DfrA and DfrB.

**Table S5** – List of accession numbers for chromosomal and mobile sequences containing DfrA/FolA-encoding genes used in this work. The species (chromosomal) or *dfrA* index (mobile), the nucleotide and protein accession numbers are provided.

**Table S6** – Protein identity values between a set of non-redundant (<90% identity) previously reported DHFR proteins resulting from pairwise alignments using the Needleman-Wunsch algorithm.

**Table S7** – %GC content for previously reported *dfrA* genes and their host genome.

**Table S8** – Analysis of %GC content among antibiotic resistance genes (ARG), the mobile genetic elements (MGE) encoding them and the species harboring the MGE. The table lists the %GC content of *dfrA/sul* homologs encoded in complete mobile element assemblies, %GC content of mobile elements and that of the host genome. Accession numbers for complete assemblies, Sul and DfrA proteins are provided.

**Table S9** – Non-redundant DfrA and Sul sequences (pairwise percent identity lower than 90%).

**Table S10** – Pairwise percent amino acid identity between aligned representative mobile DfrA protein sequences and the sequence of the chromosomal FolA protein for the species harboring the mobile genetic element encoding the *dfrA* gene.

**Table S11** – Pairwise percent amino acid identity and %GC difference between aligned representative DfrA protein sequences harbored by mobile genetic elements of *E. coli* and *K. pneumoniae*.

**Table S12** – Maximum-likelihood estimates for ancestral states in the phylogenetic tree of FolA/DfrA homologs reported in Figure 1.

**Table S13** – Pair-wise percent identities among all species in the genera under study and with the putative chromosomal origin for different *dfrA* genes. The results (p-value) of the Mann-Whitney U test comparing intra-genus pair-wise identities versus the chromosomal origin are reported for each *dfrA* gene.

**Table S14** – Percent identity values between sequences of novel mobile DHFR homologs reported here and all previously reported DHFR proteins. Percent identity values are derived from pairwise alignments using the Needleman-Wunsch algorithm.

**Table S15** – Comparison of %GC content between *dfrA* genes and putative chromosomal *folA* origins. The average coding %GC content of donor genomes is also provided.

**Table S16** – Protein sequence identity values from pairwise alignments among cryptic plasmid- and phage-encoded DHFR homologs.

## References

1. Letunic I, Bork P. Interactive Tree Of Life (iTOL) v4: recent updates and new developments. Nucleic Acids Res 2019;47:W256–W259.

2. Carlet J, Rambaud C, Pulcini C. Save Antibiotics: a call for action of the World Alliance Against Antibiotic Resistance (WAAAR). BMC Infectious Diseases 2014;14:436.

3. Rossolini GM, Arena F, Pecile P, Pollini S. Update on the antibiotic resistance crisis. Curr Opin Pharmacol 2014;18:56–60.

4. Davies J, Davies D. Origins and Evolution of Antibiotic Resistance. Microbiol Mol Biol Rev 2010;74:417–433.

5. Baym M, Lieberman TD, Kelsic ED, Chait R, Gross R, et al. Spatiotemporal microbial evolution on antibiotic landscapes. Science 2016;353:1147–1151.

6. Hegreness M, Shoresh N, Damian D, Hartl D, Kishony R. Accelerated evolution of resistance in multidrug environments. PNAS 2008;105:13977–13981.

7. Aminov RI, Mackie RI. Evolution and ecology of antibiotic resistance genes. FEMS Microbiology Letters 2007;271:147–161.

8. Sengupta S, Chattopadhyay MK, Grossart H-P. The multifaceted roles of antibiotics and antibiotic resistance in nature. Front Microbiol; 4. Epub ahead of print 2013. DOI: 10.3389/fmicb.2013.00047.

9. Kmeid JG, Youssef MM, Kanafani ZA, Kanj SS. Combination therapy for Gram-negative bacteria: what is the evidence? Expert Rev Anti Infect Ther 2013;11:1355–1362.

10. Williams K. The introduction of ‘chemotherapy’ using arsphenamine – the first magic bullet. J R Soc Med 2009;102:343–348.

11. Aminov RI. A Brief History of the Antibiotic Era: Lessons Learned and Challenges for the Future. Front Microbiol; 1. Epub ahead of print 8 December 2010. DOI: 10.3389/fmicb.2010.00134.

12. Masters PA, O’Bryan TA, Zurlo J, Miller DQ, Joshi N. Trimethoprim-Sulfamethoxazole Revisited. Arch Intern Med 2003;163:402–410.

13. Sköld O. Resistance to trimethoprim and sulfonamides. Vet Res 2001;32:261–273.

14. Quinlivan EP, McPartlin J, Weir DG, Scott J. Mechanism of the antimicrobial drug trimethoprim revisited. FASEB J 2000;14:2519–2524.

15. Hitchings GH. Mechanism of Action of Trimethoprim-Sulfamethoxazole—I. J Infect Dis 1973;128:S433–S436.

16. Landy M, Larkum NW, Oswald EJ, Streightoff F. Increased synthesis of p-aminobenzoic acid associated with the development of sulfonamide resistance in Staphylococcus aureus. Science 1943;97:265–267.

17. Huovinen P, Sundström L, Swedberg G, Sköld O. Trimethoprim and sulfonamide resistance. Antimicrob Agents Chemother 1995;39:279–289.

18. Flensburg J, Sköld O. Massive overproduction of dihydrofolate reductase in bacteria as a response to the use of trimethoprim. Eur J Biochem 1987;162:473–476.

19. Shin HW, Lim J, Kim S, Kim J, Kwon GC, et al. Characterization of trimethoprim-sulfamethoxazole resistance genes and their relatedness to class 1 integron and insertion sequence common region in gram-negative bacilli. J Microbiol Biotechnol 2015;25:137–142.

20. Rådström P, Swedberg G, Sköld O. Genetic analyses of sulfonamide resistance and its dissemination in gram-negative bacteria illustrate new aspects of R plasmid evolution. Antimicrob Agents Chemother 1991;35:1840–1848.

21. Perreten V, Boerlin P. A New Sulfonamide Resistance Gene (sul3) in Escherichia coli Is Widespread in the Pig Population of Switzerland. Antimicrob Agents Chemother 2003;47:1169–1172.

22. Tagg KA, Francois Watkins L, Moore MD, Bennett C, Joung YJ, et al. Novel trimethoprim resistance gene dfrA34 identified in Salmonella Heidelberg in the USA. J Antimicrob Chemother 2019;74:38–41.

23. White PA, Rawlinson WD. Current status of the aadA and dfr gene cassette families. J Antimicrob Chemother 2001;47:495–496.

24. Toulouse JL, Edens TJ, Alejaldre L, Manges AR, Pelletier JN. Integron-Associated DfrB4, a Previously Uncharacterized Member of the Trimethoprim-Resistant Dihydrofolate Reductase B Family, Is a Clinically Identified Emergent Source of Antibiotic Resistance. Antimicrob Agents Chemother; 61. Epub ahead of print 24 April 2017. DOI: 10.1128/AAC.02665-16.

25. Sánchez-Osuna M, Cortés P, Barbé J, Erill I. Origin of the Mobile Di-Hydro-Pteroate Synthase Gene Determining Sulfonamide Resistance in Clinical Isolates. Frontiers in Microbiology; 9. Epub ahead of print January 2019. DOI: 10.3389/fmicb.2018.03332.

26. Taly J-F, Magis C, Bussotti G, Chang J-M, Di Tommaso P, et al. Using the T-Coffee package to build multiple sequence alignments of protein, RNA, DNA sequences and 3D structures. Nat Protoc 2011;6:1669–1682.

27. O’Leary NA, Wright MW, Brister JR, Ciufo S, Haddad D, et al. Reference sequence (RefSeq) database at NCBI: current status, taxonomic expansion, and functional annotation. Nucleic Acids Res 2016;44:D733–745.

28. Benson DA, Cavanaugh M, Clark K, Karsch-Mizrachi I, Lipman DJ, et al. GenBank. Nucleic Acids Res 2017;45:D37–D42.

29. Holm L, Sander C. Removing near-neighbour redundancy from large protein sequence collections. Bioinformatics 1998;14:423–429.

30. Altschul SF, Madden TL, Schaffer AA, Zhang J, Zhang Z, et al. Gapped BLAST and PSI-BLAST: a new generation of protein database search programs. Nucleic acids research 1997;25:3389–402.

31. Kaushik S, Mutt E, Chellappan A, Sankaran S, Srinivasan N, et al. Improved Detection of Remote Homologues Using Cascade PSI-BLAST: Influence of Neighbouring Protein Families on Sequence Coverage. PLoS One; 8. Epub ahead of print 20 February 2013. DOI: 10.1371/journal.pone.0056449.

32. Brum JR, Ignacio-Espinoza JC, Roux S, Doulcier G, Acinas SG, et al. Patterns and ecological drivers of ocean viral communities. Science; 348. Epub ahead of print 22 May 2015. DOI: 10.1126/science.1261498.

33. Chen Q, Zobel J, Verspoor K. Benchmarks for measurement of duplicate detection methods in nucleotide databases. Database (Oxford). Epub ahead of print 8 January 2017. DOI: 10.1093/database/baw164.

34. Wallace IM, O’Sullivan O, Higgins DG, Notredame C. M-Coffee: combining multiple sequence alignment methods with T-Coffee. Nucleic Acids Res 2006;34:1692–1699.

35. Castresana J. Selection of Conserved Blocks from Multiple Alignments for Their Use in Phylogenetic Analysis. Molecular Biology and Evolution 2000;17:540–552.

36. Ronquist F, Huelsenbeck JP. MrBayes 3: Bayesian phylogenetic inference under mixed models. Bioinformatics (Oxford, England) 2003;19:1572–4.

37. Erill I. Dispersal and regulation of an adaptive mutagenesis cassette in the bacteria domain. Nucleic Acids Research 2006;34:66–77.

38. Rambaut A, Drummond AJ, Xie D, Baele G, Suchard MA. Posterior Summarization in Bayesian Phylogenetics Using Tracer 1.7. Syst Biol 2018;67:901–904.

39. Yu G, Smith DK, Zhu H, Guan Y, Lam TT-Y. ggtree: an r package for visualization and annotation of phylogenetic trees with their covariates and other associated data. Methods in Ecology and Evolution 2017;8:28–36.

40. Pagel M, Meade A, Barker D. Bayesian estimation of ancestral character states on phylogenies. Syst Biol 2004;53:673–684.

41. Mayola A, Irazoki O, Martínez IA, Petrov D, Menolascina F, et al. RecA protein plays a role in the chemotactic response and chemoreceptor clustering of Salmonella enterica. PLoS ONE 2014;9:e105578.

42. Clinical and Laboratory Standards Institute. Methods for dilution antimicrobial susceptibility tests for bacteria that grow aerobically - Approved Standard. Sixth edition. Wayne, Pa. USA: Clinical and Laboratory Standards Institute; 2003.

43. Fling ME, Richards C. The nucleotide sequence of the trimethoprim-resistant dihydrofolate reductase gene harbored by Tn7. Nucleic Acids Res 1983;11:5147–5158.

44. Smith DR, Calvo JM. Nucleotide sequence of the E coli gene coding for dihydrofolate reductase. Nucleic Acids Res 1980;8:2255–2274.

45. Eddy SR. Accelerated Profile HMM Searches. PLOS Comput Biol 2011;7:e1002195.

46. Faltyn M, Alcock B, McArthur A. Evolution and Nomenclature of the Trimethoprim Resistant Dihydrofolate (dfr) Reductases. Epub ahead of print 10 May 2019. DOI: 10.20944/preprints201905.0137.v1.

47. van Hoek AHAM, Mevius D, Guerra B, Mullany P, Roberts AP, et al. Acquired Antibiotic Resistance Genes: An Overview. Front Microbiol; 2. Epub ahead of print 28 September 2011. DOI: 10.3389/fmicb.2011.00203.

48. Villa L, Visca P, Tosini F, Pezzella C, Carattoli A. Composite integron array generated by insertion of an ORF341-type integron within a Tn21-like element. Microb Drug Resist 2002;8:1–8.

49. Grape M, Farra A, Kronvall G, Sundström L. Integrons and gene cassettes in clinical isolates of co-trimoxazole-resistant Gram-negative bacteria. Clin Microbiol Infect 2005;11:185–192.

50. Ho PL, Wong RC, Chow KH, Que TL. Distribution of integron-associated trimethoprim-sulfamethoxazole resistance determinants among Escherichia coli from humans and food-producing animals. Lett Appl Microbiol 2009;49:627–634.

51. Volz C, Ramoni J, Beisken S, Galata V, Keller A, et al. Clinical Resistome Screening of 1,110 Escherichia coli Isolates Efficiently Recovers Diagnostically Relevant Antibiotic Resistance Biomarkers and Potential Novel Resistance Mechanisms. Front Microbiol 2019;10:1671.

52. Kumar S, Stecher G, Suleski M, Hedges SB. TimeTree: A Resource for Timelines, Timetrees, and Divergence Times. Mol Biol Evol 2017;34:1812–1819.

53. Sigala J-C, Suárez BP, Lara AR, Borgne SL, Bustos P, et al. Genomic and physiological characterization of a laboratory-isolated Acinetobacter schindleri ACE strain that quickly and efficiently catabolizes acetate. Microbiology (Reading, Engl) 2017;163:1052–1064.

54. Falagas ME, Vardakas KZ, Roussos NS. Trimethoprim/sulfamethoxazole for Acinetobacter spp.: A review of current microbiological and clinical evidence. Int J Antimicrob Agents 2015;46:231–241.

55. Pérez-Varela M, Corral J, Aranda J, Barbé J. Roles of Efflux Pumps from Different Superfamilies in the Surface-Associated Motility and Virulence of Acinetobacter baumannii ATCC 17978. Antimicrob Agents Chemother; 63. Epub ahead of print 2019. DOI: 10.1128/AAC.02190-18.

56. Vickers AA, Potter NJ, Fishwick CWG, Chopra I, O’Neill AJ. Analysis of mutational resistance to trimethoprim in Staphylococcus aureus by genetic and structural modelling techniques. J Antimicrob Chemother 2009;63:1112–1117.

57. Watson M, Liu J-W, Ollis D. Directed evolution of trimethoprim resistance in Escherichia coli. FEBS J 2007;274:2661–2671.

58. Toprak E, Veres A, Michel J-B, Chait R, Hartl DL, et al. Evolutionary paths to antibiotic resistance under dynamically sustained drug selection. Nat Genet 2012;44:101–105.

59. Swedberg G, Fermér C, Sköld O. Point Mutations in the Dihydropteroate Synthase Gene Causing Sulfonamide Resistance. In: Ayling JE, Nair MG, Baugh CM (editors). Chemistry and Biology of Pteridines and Folates. Boston, MA: Springer US. pp. 555–558.

60. Griffith EC, Wallace MJ, Wu Y, Kumar G, Gajewski S, et al. The Structural and Functional Basis for Recurring Sulfa Drug Resistance Mutations in Staphylococcus aureus Dihydropteroate Synthase. Front Microbiol; 9. Epub ahead of print 17 July 2018. DOI: 10.3389/fmicb.2018.01369.

61. Palmer AC, Kishony R. Opposing effects of target overexpression reveal drug mechanisms. Nat Commun 2014;5:4296.

62. Dale GE, Broger C, Hartman PG, Langen H, Page MG, et al. Characterization of the gene for the chromosomal dihydrofolate reductase (DHFR) of Staphylococcus epidermidis ATCC 14990: the origin of the trimethoprim-resistant S1 DHFR from Staphylococcus aureus? J Bacteriol 1995;177:2965–2970.

63. Coque TM, Singh KV, Weinstock GM, Murray BE. Characterization of dihydrofolate reductase genes from trimethoprim-susceptible and trimethoprim-resistant strains of Enterococcus faecalis. Antimicrob Agents Chemother 1999;43:141–147.

64. Barrow EW, Bourne PC, Barrow WW. Functional cloning of Bacillus anthracis dihydrofolate reductase and confirmation of natural resistance to trimethoprim. Antimicrob Agents Chemother 2004;48:4643–4649.

65. Parkhill J, Dougan G, James KD, Thomson NR, Pickard D, et al. Complete genome sequence of a multiple drug resistant Salmonella enterica serovar Typhi CT18. Nature 2001;413:848–852.

66. Ahmed SA, Awosika J, Baldwin C, Bishop-Lilly KA, Biswas B, et al. Genomic Comparison of Escherichia coli O104:H4 Isolates from 2009 and 2011 Reveals Plasmid, and Prophage Heterogeneity, Including Shiga Toxin Encoding Phage stx2. PLOS ONE 2012;7:e48228.

67. Kim M, Kim S, Ryu S. Complete Genome Sequence of Bacteriophage SSU5 Specific for Salmonella enterica serovar Typhimurium Rough Strains. Journal of Virology 2012;86:10894–10894.

68. Octavia S, Sara J, Lan R. Characterization of a large novel phage-like plasmid in Salmonella enterica serovar Typhimurium. FEMS Microbiol Lett; 362. Epub ahead of print 1 April 2015. DOI: 10.1093/femsle/fnv044.

69. Asare PT, Jeong T-Y, Ryu S, Klumpp J, Loessner MJ, et al. Putative type 1 thymidylate synthase and dihydrofolate reductase as signature genes of a novel bastille-like group of phages in the subfamily Spounavirinae. BMC Genomics 2015;16:582.

70. Muniesa M, Colomer-Lluch M, Jofre J. Could bacteriophages transfer antibiotic resistance genes from environmental bacteria to human-body associated bacterial populations? Mob Genet Elements; 3. Epub ahead of print 1 July 2013. DOI: 10.4161/mge.25847.

71. Balcazar JL. Bacteriophages as Vehicles for Antibiotic Resistance Genes in the Environment. PLoS Pathog; 10. Epub ahead of print 31 July 2014. DOI: 10.1371/journal.ppat.1004219.

72. Enault F, Briet A, Bouteille L, Roux S, Sullivan MB, et al. Phages rarely encode antibiotic resistance genes: a cautionary tale for virome analyses. ISME J 2017;11:237–247.

73. Bhattacharya B, Giri N, Mitra M, Gupta SKD. Cloning, characterization and expression analysis of nucleotide metabolism-related genes of mycobacteriophage L5. FEMS Microbiol Lett 2008;280:64–72.

74. Wittmann J, Gartemann K-H, Eichenlaub R, Dreiseikelmann B. Genomic and molecular analysis of phage CMP1 from Clavibacter michiganensis subspecies michiganensis. Bacteriophage 2011;1:6–14.

75. Huang S, Zhang S, Jiao N, Chen F. Comparative Genomic and Phylogenomic Analyses Reveal a Conserved Core Genome Shared by Estuarine and Oceanic Cyanopodoviruses. PLOS ONE 2015;10:e0142962.

76. Kozloff LM, Lute M, Crosby LK. Bacteriophage T4 virion baseplate thymidylate synthetase and dihydrofolate reductase. J Virol 1977;23:637–644.

77. Peters DL, McCutcheon JG, Stothard P, Dennis JJ. Novel Stenotrophomonas maltophilia temperate phage DLP4 is capable of lysogenic conversion. BMC Genomics 2019;20:300.

