## Supplementary Figure 1 for "Recurrent mobilization of ancestral and novel variants of the chromosomal di-hydrofolate reductase gene drives the emergence of clinical resistance to trimethoprim"

```
                 10        20        30        40        50        60        70
            ....|....|....|....|....|....|....|....|....|....|....|....|....|....|
dfrA34      -----------------MITACVA-IDSDGGFGAQGTLPWA-IPEEFAFYQEHVRG-GICIIGGRSFNDL
dfrA9       ---------------MASLNMIVA-VNKTGGIGFENQIPWH-EPEDLKHFKAVTMN-SVLIMGRKTFASL
dfrA8       ---------------MIELHAILA-ATANGCIGKDNALPWPPLKGDLARFKKLTMG-KVVIMGRKTYESL
dfrA24      ----MTYQLDVSKILSFDLEAIVA-ATENGGIGYKGDLPWR-LQGDLKRFREITQG-GIVIMGAGTYKSL
dfrA3b      ----------------MTKQAIFA-VAENLAFGLGGGLPWDTLKDDLQFFKRLTEG-TDLVMGASTYRTL
dfrA10      ----------------MNISLIFANELITRAFGNQGKLPWQFIKEDMQFFQKTTEN-SVVVMGLNTWRSL
dfrA17      ----------------MKISLISA-VSENGVIGSGPDIPWS-VKGEQLLFKALTYN-QWLLVGRKTFDSM
dfrA32      ----------------MKISLISA-VSENGVIGSGPDIPWS-AKGEQLIFKALTYN-QWLLVGRKTFDSM
dfrA7       ----------------MKISLISA-TSENGVIGNGPDIPWS-AKGEQLLFKALTYN-QWLLVGRKTFDSM
dfrA15      ----------------MKLSLMAA-ISKNGVIGNGPDIPWS-AKGEQLLFKAITYN-QWLLVGRKTFESM
dfrA15b     ----------------MKLSLMAA-ISKNGVIGNGPDIPWS-AKGEQLLFKAITYN-QWLLVGRKTFESM
dfrA1       ----------------MKLSLMVA-ISKNGVIGNGPDIPWS-AKGEQLLFKAITYN-QWLLVGRKTFESM
dfrA28      ---------------------MVA-ISKNGVIGNGPDIPWS-AKGEQLLFKAITYN-QWLLVGRKTFESM
dfrA16      ----------------MKLSLMAA-KSKNGIIGNGPDIPWS-AKGQQLLFRAIIYN-QWLLVGRKTFESM
dfrA16b     ----------------MKLSLMAA-KSKNGIIGNGPDIPWS-AKGEQLLFKAITYN-QWLLVGRKTFESM
dfrA5       ----------------MKVSLMAA-KAKNGVIGCGPHIPWS-AKGEQLLFKALTYN-QWLLVGRKTFESM
dfrA30      ----------------MKVSLMAA-RARNGVIGCGPHIPWS-AKGEQLLFKALTYN-QWLLVGRKTFESM
dfrA14      ----------------MKVSLIAA-KRKNGVIGCGPDIPWS-AKGEQLLFKALTYN-QCLLVGRKTFESM
dfrA25      ---------------------MAA-RAKNGVIGCGPDIPWS-AKGEQLLFKALTYN-QWLLVGRKTFESM
dfrA29      ---------------------MAA-RAKNGVIGCGPNIPWS-AKGEQLLFKALTYN-QWLLVGRKTFESM
dfrA27      ----------------MKISLMAA-KARNGVIGCGSDIPWN-AKGEQLLFKAITYN-QWLLVGRKTFEAM
dfrA6       ----------------MKISLMAA-VSENGVIGSGLDIPWH-VQGEQLLFKAMTYN-QWLLVGRKTFDSM
dfrA31      ----------------MKISIMAA-VSENGVIGSGLDIPWH-VQGEQLLFKAMTYN-HWLLVGRKTFDSM
dfrA13      -----------MNPESVRIYLVAA-MGANRVIGNGPDIPWK-IPGEQKIFRRLTES-KVVVMGRKTFESI
dfrA21      -----------MNPESVRIYLVAA-MGANRVIGNGPDIPWK-IPGEQKIFRRLTES-KVVVMGRKTFESI
dfrA22      -----------MNPELVRIYLVAA-MGANRVIGNGPDIPWK-IPGEQKIFRRLTEG-KVVVMGRKTFESI
dfrA33      -----------MNPESVRIYLVAA-MGANRVIGNGPDIPWN-IPGEQKIFRRLTEG-KVVVMGRKTFESI
dfrA12      -----------MNSESVRIYLVAA-MGANRVIGNGPNIPWK-IPGEQKIFRRLTEG-KVVVMGRKTFESI
dfrF        -----------------MIGLIVA-RSKNNVIGKNGNIPWK-IKGEQKQFRELTTG-NVVIMGRKSYEEI
dfrA3       ----------------MLISLIAA-LAHNNLIGKDNLIPWH-LPADLRHFKAVTLG-KPVVMGRRTFESI
dfrA26      MADEEYDPLLDDDMEDAKVAVIAA-RAQNGCIGRHGKLPWK-LPGDLKYFRERTWG-KPIIMGRKTWESL
dfrG        ----------------MKVSLIAA-MDKNRVIGKENDIPWR-IPKDWEYVKNTTKG-HPIILGRKNLESI
dfrK        ----------------MKVSLIAA-MDKNRVIGKENDIPWR-IPEDWEYVKNTTKG-YPIILGRKNLESI
dfrD        ----------------MKISLIVA-MDKKRVIGKDNDIPWR-ISSDWEYVKNTTKG-HAIILGRKNLQSI
dfrA20      --------------MGIKYSLIVA-IGKHREMGADNDLLWH-LPRDMQFFKETTTG-HAVVMGRKSWESI
S1_dhfr     ----------------MTLSIIVA-HDKQRVIGYQNQLPWH-LPNDLKHIKQLTTG-NTLVMARKTFNSI
dfrE        -----------------MLAAIWA-QDEQGVIGKEGKLPWH-LPNDLKFFKEKTIH-NTLVLGRATFEGM
dfrA18      -------------MSHPQLELIVA-VDSKLGFGKGGKIPWK-CKEDMARFTRISKEIRVCVIGKHTYTDM
dfrA19      -------------MSHPQLELIVA-VDSKLGFGKGGKIPWK-CKEDMARFTRISKEIRVCVIGKHTYTDM
dfrA23      ---------------MPTVEIIVA-VDPVGGFGRNGQIPWT-CKEDMKRFTTISKEIRVCVMGKNTYKDM
dfrB5       ------------------------------MDQGRSEVSNP-VAGQFAFPSNAAFG-----MGDRVRKKS
dfrB7       ------------------------------MDQGSNEVGNP-VAGQFSFPSNAAFS-----MGDRVRKKS
dfrB6       ------------------------------MDQGSNEVINP-VAGQFASPSNATFG-----MGDRVRKKS
dfrB1       ----------MQRVVGPHRTPRSS-QERSEMERSSNEVSNP-VAGNFVFPSNATFG-----MGDRVRKKS
dfrB2       ------------------------------MGQSSDEANAP-VAGQFALPLSATFG-----LGDRVRKKS
dfrB3       ------------------------------MDQHNNGVSTL-VAGQFALPSHATFG-----LGDRVRKKS
dfrB4       ------------------------------MNEGKNEVSTS-AAGRFAFPSNATFA-----LGDRVRKKS
```

**Figure S1 –** Multiple sequence alignment including all reported mobile DHFR proteins. DfrB protein sequences are highlighted in yellow.

**Figure S1 –** (continued).

```
                     80        90       100       110       120       130       140
         ....|....|....|....|....|....|....|....|....|....|....|....|....|....|
dfrA34   VHLSLSPKGGLYKKCLLRTTPHIVVSSSHELVYDPSIMALIEADRRHLDLYFVNTVDAAVKLAKGLGG--
dfrA9    P---------------------KVLPGRLHVVVSKTVPPTQNTDQVVYVSTYQIAVRTASLLVDKP----
dfrA8    P---------------------VKLEGRTCIVMTRQALELPGVRDAN-GAIFVNNVSDAMRFAQEES---
dfrA24   P---------------------SPLKDRINIVITKKSEISWTACY---DVRVVNSPEDALRMVGRIIDEK
dfrA3b   P--------------------LLPTNNRQFIVVSNTEEPSLNVH-----VVSPEHFKAFLSKTSR-----
dfrA10   P--------------------KMKKLGRDFIVISSTITEHEVLNN---NIQIFKSFESFLEAFRD-----
dfrA17   G----------------------VLPNRKYAVVSKNGISSSNE-----NVLVFPSIENALKELSK-----
dfrA32   G----------------------VLPNRKYAVVSKNGISGSNE-----NVLVFPSIENALQELSK-----
dfrA7    G----------------------VLPNRKYAVVSRKGISSSNE-----NVLVFPSIEIALQELSK-----
dfrA15   G----------------------ALPNRKYAVVTRSSFTSSDE-----NVLVFPSIDEALNHLKT-----
dfrA15b  G----------------------ALPNRKYAVVTRSSFTSSDE-----NVLVFPSIDEALNHLKT-----
dfrA1    G----------------------ALPNRKYAVVTRSSFTSDNE-----NVLIFPSIKDALTNLKK-----
dfrA28   G----------------------ALPNRKYAVVTRSSLTSDNE-----NVVIFPSIKDALTNLKK-----
dfrA16   G----------------------ALPNRKYAVVTRSNFSTNDE-----GVMVFSSIQDALINLEE-----
dfrA16b  G----------------------ALPNRKYAVVTRSNFSTNDE-----GVMVFSSIQDALINLQE-----
dfrA5    G----------------------ALPNRKYAVVTRSAWTADND-----NVIVFPSIEEAMYGLAE-----
dfrA30   G----------------------ALPNRKYAVVTRSAWTANND-----NVVVFPSIEEAMGGLAK-----
dfrA14   G----------------------ALPNRKYAVVTRSGWTSNDD-----NVVVFQSIEEAMDRLAE-----
dfrA25   G----------------------PLPNRKYAVVTRSNWTAANE-----NVVVFPSIDEAMGRLGE-----
dfrA29   G----------------------PLPNRKYAVVTRSNWTAANE-----NVVVFPSIDEAMGRLGE-----
dfrA27   G----------------------ALPNRKYAVVSRSGSVATND-----DVVVFPSIEAAMRELKT-----
dfrA6    G----------------------KLPNRKYAVVTRSKIISNDP-----DVVYFASVESALAYLNN-----
dfrA31   G----------------------KLPNRKYAVVTRSEMVSNDP-----DVIYFTSIESALSYLDN-----
dfrA13   G---------------------KPLPNRHTVVLSRQAGYSAPG------CAVVSTLSHVSPSTAE-----
dfrA21   G---------------------KPLPNRHTVVLSRQARYSAPG------CAVVSTLSQAIAIAAE-----
dfrA22   G---------------------KPLPNRRTVVLSRQASYSAAG------CAVVSTLSQAIAIAAE-----
dfrA33   G---------------------KPLPNRRTVVLSRQASYSAAG------CAVVSTLSQAIAIAAE-----
dfrA12   G---------------------KPLPNRHTLVISRQANYRATG------CVVVSTLSHAIALASE-----
dfrF     G---------------------HPLPNRMNIVVSTTTEYQGDN------LVSVKSLEDALLLAK------
dfrA3    G---------------------RPLPGRRNVVVSRNPQWQAEG------VEVAPSLDAALALLTD-----
dfrA26   N---------------------GALPGRTNIVVTRQQGYEAEGARVVDSIEEAISLAQSIALIEA-----
dfrG     G---------------------RALPDRRNIILTRDKGFTFNG------CEIVHSIEDVFELCKN-----
dfrK     G---------------------RALPGRRNIILTRDKGFSFNG------CEIVHSIEDVFELCNS-----
dfrD     G---------------------RALPDRRNIILTRDKNFNFKD------CEIAHSIEAAFKLCEN-----
dfrA20   PQK------------------YRPLPNRLNFVLTRDKNYSAEG------ATVIYDLKEVAQHLEG-----
S1_dhfr  G---------------------KPLPNRRNVVLTNQASFHHEG------VDVINSLDEIKELS-------
dfrE     GC--------------------RPLPNRTTIVLTSNPDYQAEG------VLVMHSVEEILAYADK-----
dfrA18   RDMQLEKDG-----AEERIKEKGILPERESFVISSTLKQEDVIG-----ATVVPDLRAVINLYEN-----
dfrA19   RDMQLEKDG-----AEERIKEKGILPERESFVISSTLKQEDVIG-----ATVVPDLRAVINLYEN-----
dfrA23   LDMQMKKEG-----AEERIKEKGILPERESYVVSSTLKPEDVIG-----ATVVPDLRAVLNQYHD-----
dfrB5    G-----------------------------------AAWQG-----------------------------
dfrB7    G-----------------------------------AAWQG-----------------------------
dfrB6    G-----------------------------------AAWQG-----------------------------
dfrB1    G-----------------------------------AAWQG-----------------------------
dfrB2    G-----------------------------------AAWQG-----------------------------
dfrB3    G-----------------------------------AAWQG-----------------------------
dfrB4    G-----------------------------------AAWQG-----------------------------
```

**Figure S1 –** (continued).

```
                 150       160       170       180       190       200       210
            ....|....|....|....|....|....|....|....|....|....|....|....|....|....|
dfrA34      ---MHANKDIHFIGGKRIYDAGLD--YCDEVYTSILPAVYLN-CDTFFPVEKLSRMFTPELYKTIPNQ--
dfrA9       -----EYSQIFVIGGKSAYENLAA--YVDKLYLTRVQLNTQQ--DTELDLSLFKSWKLVSEVPTITEN--
dfrA8       -----VGDVAYVIGGAEIFKRLAL--MITQIELTFVKRLYEGD-TYVDLAEMVKDYEQNGMEEHDLHT--
dfrA24      EEQGRDRPRVFVIGGASIYQALMP--FVSTLHWTEVHVEQLPE-EIGLDTYIEDFLSLRGTSTPKRKS--
dfrA3b      --------NLTIIGGSSLLTVDILS-KMDKIIMTTVYGSFDA--DVYLPTEVVSYVTGKASNATLFNN--
dfrA10      -----TTKPINVIGGVGLLSEAIE--HASTVYMSSIHMVKPVHADVYVPVELMNKLYSDFKYPENILWVG
dfrA17      -----VTDHVYVSGGGQIYNSLIE--KADIIHLSTVHVEVEG--DIKFPIM-PENFNLVFEQFFMSNI--
dfrA32      -----ITDHVYISGGGQIYESLIE--KADIIHLSTIHVEVEG--DIKFPIL-PEGFNLVFEQFFVSNI--
dfrA7       -----ITDHLYVSGGGQIYNSLIE--KADIIHLSTVHVEVEG--DINFPKI-PENFNLVFEQFFLSNI--
dfrA15      -----ITDHVIVSGGGEIYKSLID--KVDTLHISTIDIEPEG--DVYFPEI-PSSFRPVFSQDFVSNI--
dfrA15b     -----ITDHVIVSGGGEIYKSLID--KADTLHISTIDIEPEG--DVYFPEI-PGSFRPVFSQDFVSNI--
dfrA1       -----ITDHVIVSGGGEIYKSLID--QVDTLHISTIDIEPEG--DVYFPEI-PSNFRPVFTQDFASNI--
dfrA28      -----ITDHVIVSGGGEIYKSPID--QVDTLHISTIDIEPEG--DVYFPES-PAILG-QFYPRLRSNI--
dfrA16      -----ITDHVIVSGGGEIYKSLIS--KVDTLHISTVDIERDG--DIVFPEI-PDTFKLVFEQDFESNI--
dfrA16b     -----ITDHVIVSGGGEIYKSLIS--KVDTLHISTVDIERDG--DIVFPEI-PDTFKLVFEQDFESNI--
dfrA5       -----LTDHVIVSGGGEIYRETLP--MASTLHISTIDIEPEG--DVFFPNI-PNTFEVVFEQHFSSNI--
dfrA30      -----LNGHVIVSGGGEIYRETLP--MASTLHVSTIDIEPEG--DVFFPNI-PNFFEVVFEQHFSSNI--
dfrA14      -----FTGHVIVSGGGEIYRETLP--MASTLHLSTIDIEPEG--DVFFPSI-PNTFEVVFEQHFTSNI--
dfrA25      -----ITDHVIVAGGGEIYHETIP--MASTLHVSTIDVEPEG--DVFFPNI-PGKFDVVFEQQFTSNI--
dfrA29      -----ITDHVIVAGGGEIYHETIP--MASTLHVSTIDVEPEG--DVFFPNI-PGKFDVVFEQQFTSNI--
dfrA27      -----LTNHVVVSGGGEIYKSLIA--HADTLHISTIDSEPEG--NVFFPEI-PKEFNVVFEQELHSNI--
dfrA6       -----ATAHIFVSGGGEIYKALID--QADVIHLSVIHKHISG--DVFFPPV-PQGFKQTFEQSFSSNI--
dfrA31      -----TTTHVFVSGGGEIYKALIE--QADVIHLSVIHKHISG--DVFFPSV-PQSFKQTFEQSFSSNI--
dfrA13      -----HGKELYVARGAEVYALALP--HANGVFLSEVHQTFE-G-DAFFPVLNAAEFEVVSSETIQGTI--
dfrA21      -----HGKELYVAGGAEVYALALP--HANGVFLSEVHQTFE-G-DAFFPVLNAAEFEVVSSETIQGTI--
dfrA22      -----HGKELYVAGGAEVYALALP--RADGVFLSEVHQTFE-G-DAFFPVLDEAEFEVVSAETVQATI--
dfrA33      -----HGKELYVAGGAEVYALALP--RADGVFLSEVHQTFE-G-DAFFPALDAAEFDVVSAETVQATI--
dfrA12      -----LGNELYVAGGAEIYTLALP--HAHGVFLSEVHQTFE-G-DAFFPMLNETEFELVSTETIQAVI--
dfrF        ------GRDVYISGGYGLFKEALQ--IVDKMYITEVDLNIEDG-DTFFPEFDINDFEVLIGETLGEEV--
dfrA3       -----C-EEAMIIGGGQLYAEALP--RADRLYLTYIDAQLNG--DTHFPDYLSLGWQELERSTHPADD--
dfrA26      -----V-DEIMVLGGGEIYTQALP--QADILYLTEVHASVDG--DAFFPDVDLSQYQETQRQDFEPSG--
dfrG        -----E-EEIFIFGGEQIYNLFFP--YVEKMYITKIHHEFEG--DTFFPEVNYEEWNEVFAQKGIKND--
dfrK        -----E-EEIFIFGGEQIYNLFLP--YVEKMYITKIHYEFEG--DTFFPEVNYEEWNEVSVTQGITNE--
dfrD        -----E-EEVFIFGGEQIYVMFLP--YVEKMYVTKIHHEFEG--DTFFPVVNFDDWKEVSVEKGIKDE--
dfrA20      -----KNLTCFIIGGAQIYQLALETGLLNEMYVTQVHNTFEEA-DTFFPFVNWGEWEEEDILEQDKDE--
S1_dhfr     -------GHVFIFGGQTLYEAMID--QVDDMYITVIDGKFQG--DTFFPPYTFENWEVESSVEGQLDE--
dfrE        -----YEGVTVIGGGSVVFKELIP--ACDVLYRTMIHETFEG--DTFFPEIDWSVWEKVATVPGVVDE--
dfrA18      -----TDQRIAVIGGEKLYIQALS--SATKLHMTIIPREFDCD-RFIPVDPIQNNFHIDSSASETVEAT-
dfrA19      -----TDQRIAVIGGEKLYIQALS--SATKLHMTIIPREFDCD-RFIPVDPIQNNFHIDSSASETVEAT-
dfrA23      -----SDQRIAVIGGEKLYVQALA--SATKVHMTVMHKPYNCD-RTLPMSYIDKKF-VAGQGSITIQTA-
dfrB5       ----------QIVG---WYCTKLT----PEGYAVESEAHPGS--VQIYPVAALERIN-------------
dfrB7       ----------QIVG---WYCTKLT----PEGYAVESEAHPGS--VQIYPVAALERINGVQG---------
dfrB6       ----------QIVG---WYSTKLT----PEGYAVESEAHPGS--VQIYPVAALERVN-------------
dfrB1       ----------QIVG---WYCTNLT----PEGYAVESEAHPGS--VQIYPVAALERIN-------------
dfrB2       ----------QVVG---WYCTKLT----PEGYAVESESHPGS--VQIYPVAALERVA-------------
dfrB3       ----------QVVG---WYCTKLT----PEGYAVESESHPGS--VQIYPVAALERVA-------------
dfrB4       ----------RIVG---WYCTTLT----PEGYAVESESHPGS--VQIYPMTALERVA-------------
```

**Figure S1 –** (continued).

```
                    220       230       240
          ....|....|....|....|....|....|....
dfrA34    ----VHADIPVIKWTRKRA---------------
dfrA9     -KTKLIFQIWINPNPISEEPTC------------
dfrA8     ---YFTYRKKELTE--------------------
dfrA24    ---NLVLPPTPTTP--------------------
dfrA3b    --SDAKMAVYYG----------------------
dfrA10    DPIDSVYSLSIDKFVRPASLVGVPNDINT-----
dfrA17    ---NYTYQIWKKG---------------------
dfrA32    ---NYTYQIWKKG---------------------
dfrA7     ---NYTYQIWKKG---------------------
dfrA15    ---NYSYQIWQKG---------------------
dfrA15b   ---NYSYQIWQKG---------------------
dfrA1     ---NYSYQIWQKG---------------------
dfrA28    ---NYSYQIWQRVQCSTIRNCPLCTKRQVASLAG
dfrA16    ---NYCYQIWQKS---------------------
dfrA16b   ---NYCYQIWQKS---------------------
dfrA5     ---NYCYQIWQKG---------------------
dfrA30    ---NYCYQIWKKG---------------------
dfrA14    ---NYCYQIWKKG---------------------
dfrA25    ---NYCYQIWQKG---------------------
dfrA29    ---NYCYQIWQKG---------------------
dfrA27    ---NYRYQIWQRG---------------------
dfrA6     ---DYTYQIWAKG---------------------
dfrA31    ---DYTYQIWAKG---------------------
dfrA13    ---TYTHSVYARRNG-------------------
dfrA21    ---TYTHSVYARRNG-------------------
dfrA22    ---TYTHSVYARRNG-------------------
dfrA33    ---TYTHSVYARRNG-------------------
dfrA12    ---PYTHSVYARRNG-------------------
dfrF      ---KYTRTFYVRKNELSRFWI-------------
dfrA3     ---KNSYACEFVTLSRQR----------------
dfrA26    ---GNPYPFSFVVYQRT-----------------
dfrG      ---KNPYNYYFHVYERKNLLS-------------
dfrK      ---KNPYTYYFHIYERKAS---------------
dfrD      ---KNPYDYYFHIYERIR----------------
dfrA20    ---KHLYSFNIKKFTR------------------
S1_dhfr   ---KNTIPHTFLHLVRRKGK--------------
dfrE      ---KNLYAHDYETYHR-NDK--------------
dfrA18    -VDETQERIHFATYVRNNQ---------------
dfrA19    -VDETQERIHFATYVRNNQ---------------
dfrA23    -VDGETHPVKFITYERARP---------------
dfrB5     ----------------------------------
dfrB7     ----------------------------------
dfrB6     ----------------------------------
dfrB1     ----------------------------------
dfrB2     ----------------------------------
dfrB3     ----------------------------------
dfrB4     ----------------------------------
```
