## Supplementary Figure 2 for "Recurrent mobilization of ancestral and novel variants of the chromosomal di-hydrofolate reductase gene drives the emergence of clinical resistance to trimethoprim"

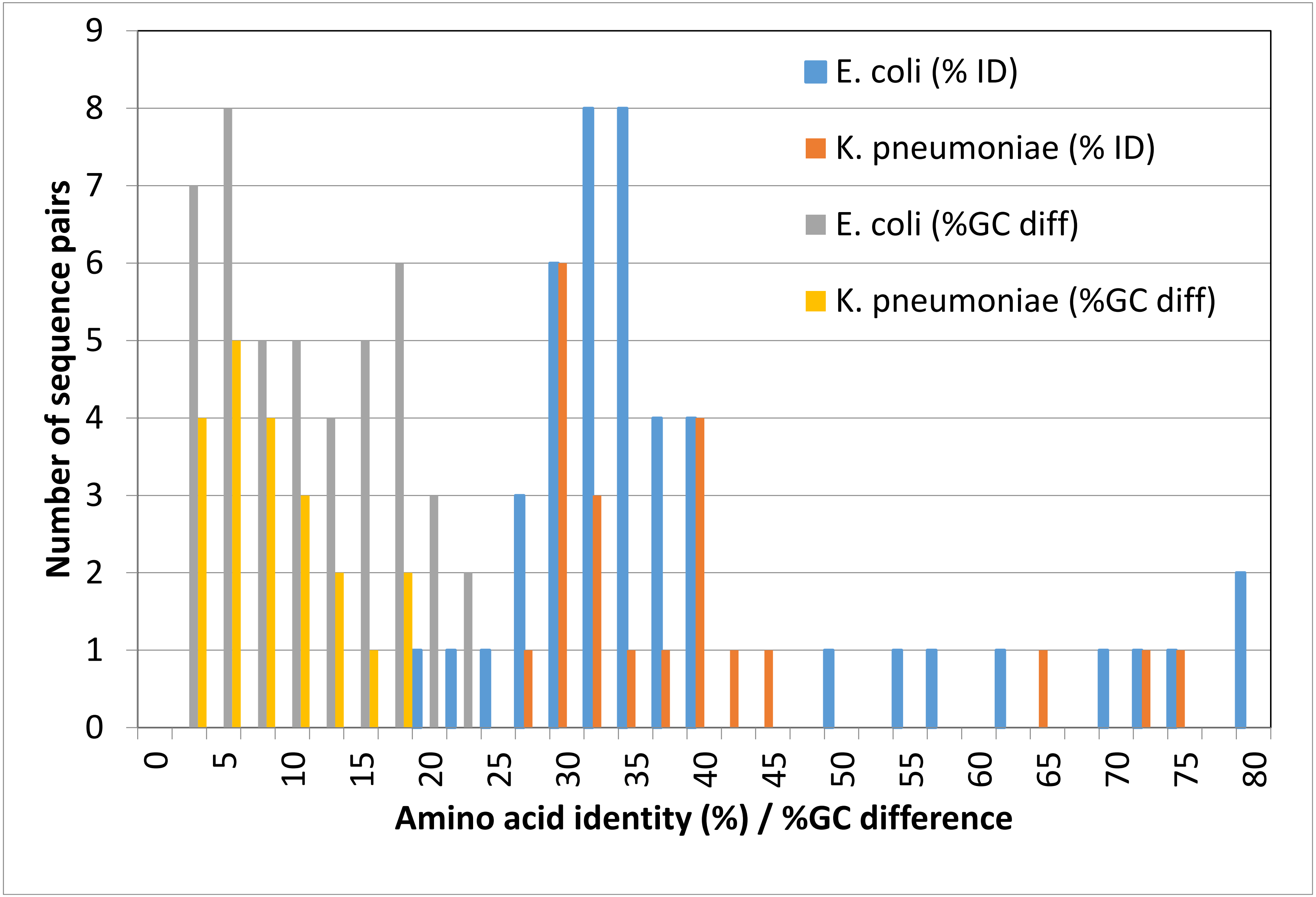


**Figure S2 –** Pairwise percent amino acid identity and %GC difference between aligned representative DfrA protein sequences harbored by mobile genetic elements of *E. coli* and *K. pneumoniae*.
