## Supplementary Figure 3 for "Recurrent mobilization of ancestral and novel variants of the chromosomal di-hydrofolate reductase gene drives the emergence of clinical resistance to trimethoprim"

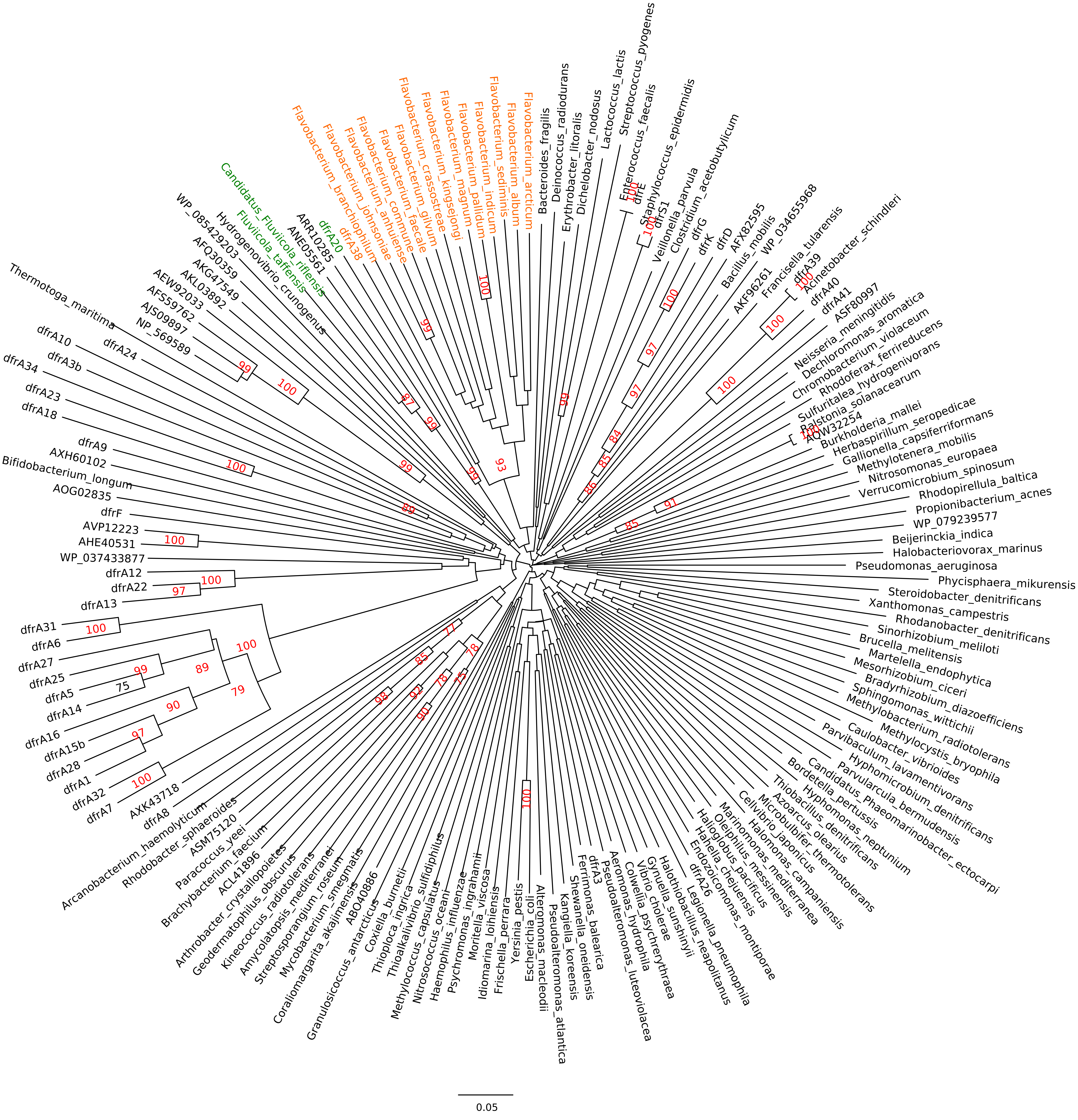


**Figure S3 –** Unrooted Neighbor-Joining tree of DHFR protein sequences. Branch support values are provided as the total number of 1000 bootstrap pseudo-replicates in which the branching was observed. Support values are only shown for branches with at least 75% bootstrap support. The DfrA20, DfrA38, *Fluviicola* and *Flavobacterium* DHFR protein sequences are highlighted.
