## Supplementary Figure 4 for "Recurrent mobilization of ancestral and novel variants of the chromosomal di-hydrofolate reductase gene drives the emergence of clinical resistance to trimethoprim"

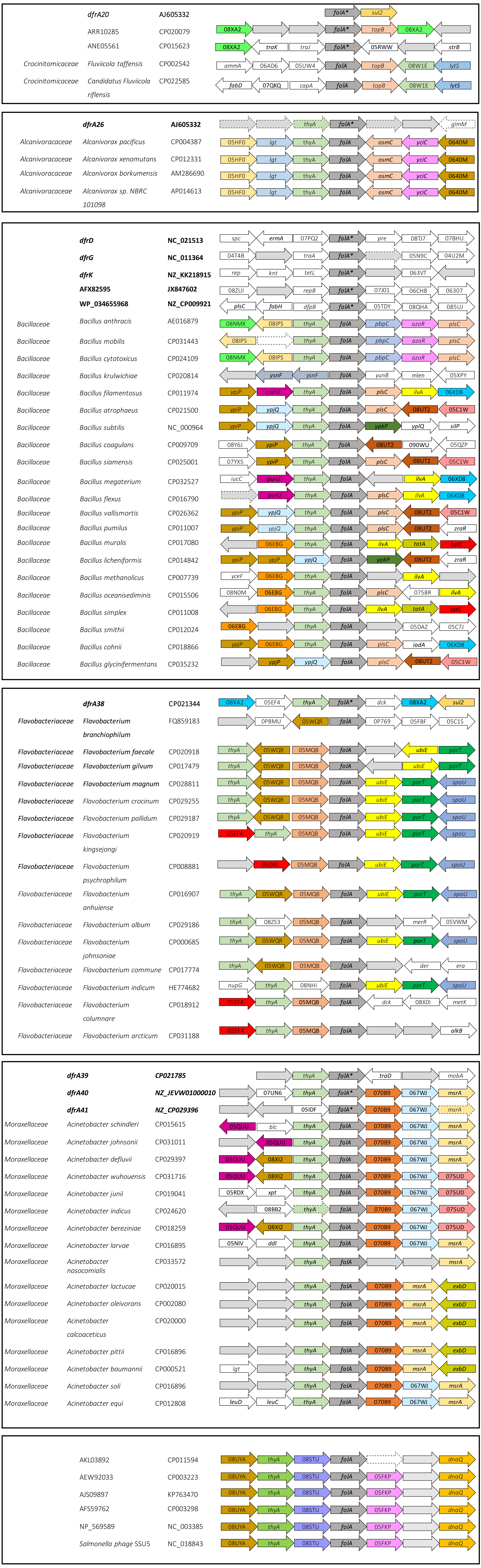


**Figure S4 –** Schematic representation of the genetic environment of mobile DHFR genes, their putative chromosomal origin and one representative complete genome assembly for each species within the corresponding genus. Arrow boxes indicate coding regions (discontinued arrows pinpoint pseudogenes). When available, gene names or NOG identifiers are provided and color coded.
