## Supplementary Figure 5 for "Recurrent mobilization of ancestral and novel variants of the chromosomal di-hydrofolate reductase gene drives the emergence of clinical resistance to trimethoprim"

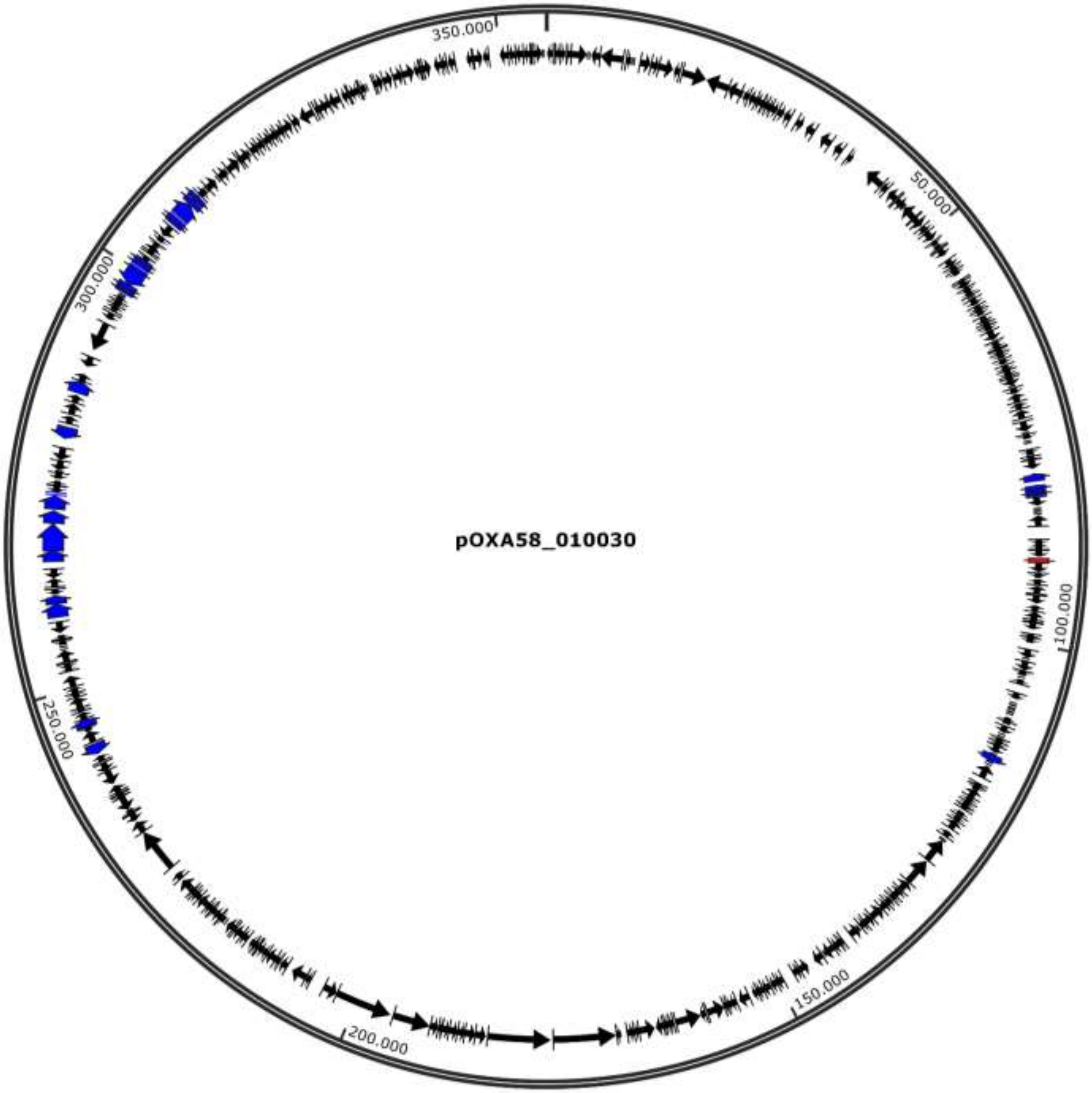


**Figure S5 –** Graphical overview of *Acinetobacter defluvii* plasmid pOXA58_010030 with *dfrA41* (red arrow box) and other resistance determinants represented as blue arrow boxes. This figure was constructed using SnapGene Viewer.
